# KANK2 at focal adhesion regulates their maintenance and dynamics, while at fibrillar adhesions it influences cell migration via microtubule-dependent mechanism

**DOI:** 10.1101/2025.02.24.639831

**Authors:** Nikolina Stojanović, Anja Rac, Marija Lončarić, Ana Tadijan, Mladen Paradžik, Marta Acman, Jonathan D. Humphries, Martin J. Humphries, Andreja Ambriović-Ristov

**Author notes:** equal contribution (Nikolina Stojanović, Anja Rac).

## Abstract

**Background:** Integrins form focal adhesions (FAs) at the cell edge and fibrillar adhesions (FBs) located centrally. Talin 1 is essential to FAs, while talin2 is found in FAs and FBs. KANK (kidney ankyrin repeat-containing) family proteins regulate adhesion dynamics and are recruited to adhesions through interaction with talins. Previously, we showed in MDA-MB-435S melanoma cells that KANK2 is part of integrin αVβ5 FAs, that its interaction with talin2 regulates actin-microtubule (MT) crosstalk and that KANK2 knockdown mimics the effect of integrin αV or β5 knockdown by reducing cell migration. Here, in another melanoma cell line RPMI-7951 we observed that KANK2 is part of FAs and FBs and that KANK2 knockdown increases cell migration. Therefore, we analyse integrin adhesion complexes in RPMI-7951 cells, explore the localization and role of KANK2 in FAs and FBs.

**Methods:** Knockdown in human melanoma RPMI-7951 cells was achieved by transfection with gene-specific siRNAs. Integrin adhesion complexes were isolated and analysed by mass spectrometry. Immunofluorescence analysis, live cell imaging and the proximity ligation assays were done using confocal microscopy. Cell migration was assessed using Transwell Cell Culture Inserts. MTT assays were performed to determine cell sensitivity to paclitaxel. Data were statistically evaluated using one-way or two-way ANOVA or unpaired Student’s t-tests in GraphPad Software.

**Results:** We demonstrate that RPMI-7951 melanoma cells use integrin αVβ5 FAs and integrin α5β1 FBs for adhesion, and that KANK2 is part of both structures. KANK2 is predominantly in proximity to talin1 at the cell edge (FAs) and in proximity to talin2 in the cell center (FBs). KANK2 in FAs functionally interacts with talin1 to maintain FAs, and with talin2 to regulate their dynamics. KANK2 is a component of FBs, and its knockdown mimics integrin α5 knockdown by increasing MT-dependent cell migration.

**Conclusions:** Our study reveals the distinct roles of KANK2 in FAs and FBs. We show that KANK2 is a component of FBs, linking them to MTs and promoting their stabilisation. Loss of integrin α5 or KANK2 from FBs increases cell migration, a process that relies on the MT cytoskeleton.

**Plain English summary:** Cells adhere to their surroundings using specialized structures called adhesions, which are formed by different types of integrins and have different cell localisations: focal adhesions at the cell edge and fibrillar adhesions in the cell center. Talin1 is essential for focal adhesions, while talin2 is present in focal and fibrillar adhesions. KANK (kidney ankyrin repeat-containing) family proteins regulate adhesion dynamics by connecting talins to the microtubule cytoskeleton. Our study explored the different roles of KANK2 in focal or fibrillar adhesions, by focusing on its interactions with integrins and talins. We show that in RPMI-7951 cells, which form αVβ5 focal and α5β1 fibrillar adhesions, KANK2 is present in both of these adhesions. KANK2 supports αVβ5 focal adhesion maintenance via talin1 and regulates their dynamics via talin2. KANK2 is also a component of α5β1 fibrillar adhesions, and its depletion mimics integrin α5 knockdown and enhances cell migration, a process dependent on the microtubule cytoskeleton. In conclusion, our study demonstrates that KANK2 plays distinct roles in different adhesion structures, contributing to both focal and fibrillar adhesion function.

## INTRODUCTION

Integrins are transmembrane cell surface proteins consisting of α and β subunits that combine to form 24 distinct heterodimers, enabling cell-to-cell or cell-to-extracellular matrix (ECM) connections [1]. Integrin heterodimer binding of ECM ligands results in the formation of several functionally and structurally different classes of integrin-based adhesions. These include short-lived peripheral nascent adhesions, which mature into focal adhesions (FAs) and further, centrally located, elongated fibrillar adhesions (FBs), which all serve as a link to the actin cytoskeleton [2]. Additionally, there are hemidesmosomes, which link to intermediate filaments [3], and reticular adhesions (RAs), which are not associated to the cytoskeleton [4].

Integrins are involved in the metastatic cascade and have been found to be responsible for intrinsic and acquired therapy resistance. Therefore, targeting integrins, or more specifically the signal transduction *via* integrin adhesion complexes (IACs), is a promising therapeutic opportunity to reduce tumor metastasis and overcome therapeutic resistance [5]. However, despite highly encouraging preclinical data, targeting IACs in clinical trials has so far failed [6, 7]. Therefore, further work is required not only to understand the role of different types of adhesions, their turnover and the relationships between adhesion structures, but also to unlock their therapeutic potential.

FAs are the most extensively studied IACs that convert extracellular chemical and mechanical cues into biochemical signals [8, 9]. In culture, cells form FAs in order to adhere to their substrate. FAs link integrin heterodimers bound to ECM outside the cell and the actomyosin and microtubule (MT) network inside the cell, acting *via* a range of adaptor proteins [10, 11]. FBs develop from mature FAs and have a role in promoting fibronectin remodeling [12]. They are larger than FAs, with a dissimilar composition, and are located in the center of the migrating cell [13, 14]. There are multiple integrin heterodimers commonly associated with FAs (e.g. αVβ3, αVβ5, α5β1, α3β1 and others) while integrin α5β1 is primarily involved in forming FBs [15]. In contrast to FAs and FBs, RAs represent a specialized adhesions mediated by integrin αVβ5. They are distinct from other adhesion types due to their lack of cytoskeletal connections and their high stability, making them uniquely suited for signaling and maintaining cell-ECM adhesion in situations where cytoskeletal connections are not required, such as cell division [4, 16–20].

Talins are key cytoplasmic mechanosensitive proteins mediating binding of integrin adhesions to the ECM [21]. There are two different isoforms of talins, talin1 and talin2. Although talin1 and talin2 share 76% protein sequence identity (88% similarity), they are not functionally redundant, and the differences between the two isoforms are not fully understood [22]. Talin1 forms the core of IAC, increasing the affinity of integrin for ligands (integrin activation) and mediating the mechanosensitive link to actin [23, 24], while talin2 has been relatively less studied. Talins also coordinate MT recruitment to adhesion sites *via* interaction with KANK (kidney ankyrin repeat-containing) proteins, members of the cortical microtubule stabilizing complex (CMSC) [21, 25–28].

KANK family proteins are evolutionarily conserved and consist of four paralogs (KANK1–4) due to gene expansion and diversification. KANK1 (around 170 kDa) is mainly found in epithelial, while KANK2 (around 120 kDa) in mesenchymal cells [29]. KANK1 and 2 localize to the rims of mature integrin-containing FAs, termed the FA belt and adjacent regions. The CMSC, recruited upon talin-KANK2 interaction, stabilizes MTs in the vicinity of adhesions and regulates actin-MT crosstalk [25, 30]. The uncoupling of FAs and CMSCs leads to the release of the RhoA GEF ARHGEF2 / GEF-H1 from MTs that in turn results in increased RhoA-mediated actomyosin contractility, reinforcement of the stress fiber-associated FAs and decreased FA turnover [31]. Other possible localization of KANK2, with no reports indicating similar localization for KANK1, is in centrally positioned FBs. KANK2 in FAs binds talin in the vicinity of F-actin binding site 2 (ABS2) and destabilizes the talin-F-actin linkage that leads to slippage of integrin ligand bonds and adhesion sliding in the centrally positioned adhesions [27].

A discrepancy in how KANK proteins affect cell migration and sensitivity to antitumor agents has been observed (reviewed in [32]). It is also not clear which talin isoform binds which KANK paralogue within the cell. While in HeLa cells Bouchet et al. [25] showed that talin1 binds KANK1, we have recently showed that KANK2 and talin2 functionally interact in the melanoma cell line MDA-MB-435S thus enabling the actin-MT crosstalk important for both sensitivity to MT poisons and cell migration [28].

Here, we present an analysis of IACs in the melanoma cell line RPMI-7951 cultivated in long-term culture. Our study explores the localization and roles of KANK2. We demonstrate that KANK2 is part of integrin αVβ5 FAs and α5β1 FBs. In FAs, KANK2 functionally interacts (i) with talin1 to maintain FAs, and (ii) with talin2 to regulate their dynamics. Our data also reveal that KANK2 localized within FBs regulates cell migration *via* MTs.

## MATERIALS AND METHODS

### Cell Culture

The human melanoma cell line RPMI-7951 was obtained from the American Type Culture Collection (ATCC). Cells were grown on uncoated cell culture plates in DMEM (Invitrogen) supplemented with 10% (v/v) FBS (Invitrogen) (DMEM-FBS) at 37°C with 5% CO2 (v/v) in a humidified atmosphere. RPMI-7951-EB3 cell populations with fluorescently labelled end-binding protein 3 (EB3) were generated by stable transfection of Dendra2-EB3-7 plasmid containing the photoconvertible fluorescent protein Dendra2 fused with EB3 (EB3-Dendra2) (Addgene plasmid #57715) using Lipofectamine 2000 (Thermo Fisher Scientific) and selection with 1.2 mg/mL geneticin (Sigma-Aldrich).

### Transient siRNA Transfection

For transient siRNA transfection, cells (3.7 × 10^4^, 5 × 10^5^ or 1 × 10^6^) were plated in 24-well, 6-well cell culture plates or 10 cm diameter Petri dishes to achieve 60–80% confluence after 24 hours. Cells were transfected 24 hours later, using Lipofectamine RNAiMax (13778150, Thermo Fisher Scientific), with 25 nM of control non-specific (Silencer™ Select Negative Control No. 1 siRNA, Ambion; si(-))), gene-specific siRNA for ITGAV (s7568, Ambion; si(αV)), ITGA5 (s7547, Ambion; si(α5)), ITGB5 (s7591, Ambion; si(β5)) or gene-specific siRNA for KANK2 (target sequence: ATGTCAACGTGCAAGATGA; si(KANK2)), TLN1 (target sequence: TGAATGTCCTGTCAACTGCTG; si(talin1)) or TLN2 (target sequence: TTTCGTTTTCATCTACTCCTT; si(talin2)) [28], all purchased from Sigma. Knockdown was validated by immunofluorescence using specific antibodies and matched labelled secondary antibodies (See Supplementary Table S1, Additional File 1).

### Isolation of IACs

IACs were isolated from cells cultivated for 72 hours as described previously [28, 33, 34]. Briefly, cells were washed with DMEM-HEPES and incubated with Wang and Richard’s reagent (DTBP, 6 mM, Thermo Fisher Scientific) for 15 minutes. DTBP was quenched with 0.03 M Tris-HCl (pH 8) and cells were lysed using modified RIPA buffer (50 mM Tris-HCl, pH 7.6; 150 mM NaCl; 5 mM disodium EDTA, pH 8; 1% (w/v) Triton X-100, 0.5% (w/v) SDS, 1% (w/v) sodium deoxycholate). Cell bodies were removed by high-pressure washing with tap water for 10 seconds and remaining adhesion complexes were collected by scraping into adhesion recovery solution (125 mM Tris-HCl, pH 6.8; 1% (w/v) SDS; 150 mM dithiothreitol). Samples containing isolated IACs were acetone-precipitated and further processed for Mass Spectrometry (MS) or Western Blotting [35].

### Sample Preparation for Mass Spectrometry and Data Analysis

Samples (prepared in triplicates) were processed as previously described [33, 34] and analysed using an UltiMate R 3000 Rapid Separation LC (RSLC, United States) coupled to an Orbitrap Elite mass detector (Thermo Fisher Scientific, United States) with electrospray ionization. Peptide mixtures were eluted over 44 min using a gradient of 92% of solution A (0.1% formic acid in water) and 8% up to 33% of solution B (0.1% formic acid in acetonitrile). Solvent flow was set to 300 nL per minute. Protein identification was performed by searching data against the human SwissProt database (version 2018_01) using Mascot (Matrix science, version 2.5.1). Fragment ion tolerance was set to 0.50 Da, while parent ion tolerance was 5 PPM. Scaffold (Proteome software) was used to further refine protein identification. Protein (99.9%) and peptide (95%) probabilities were assigned using the Protein Prophet algorithm [36] as incorporated by Scaffold including a minimum of four unique peptides per each protein. Data have been deposited in the ProteomeXchange Consortium [37] (dataset ID: PXD064756) via the PRIDE repository [38]).

### Protein-protein Interaction Network Formation, Functional Enrichment, Gene Ontology Analysis, and MS Data Visualization

Human protein–protein interactions (PPIs) were loaded from STRING database, using stringApp (confidence score cut-off = 0.40, maximum additional interactors = 0) [39] for Cytoscape software (version 3.7.1) [40]. Functional annotation was performed using the Database for annotation, visualization and integrated discovery (DAVID), version 6.8 [41, 42] and Panther GO database [43]. Functional enrichment analysis was carried out using the DAVID_CC subontology list (Benjamini–Hochberg corrected P-value < 0.05, EASE score < 0.1, at least four identified proteins). To summarize gene ontology (GO) terms and visualize them in a similarity-based space, the REVIGO tool was used with the following parameters: comparison of corrected P-values related to GO terms, allowed similarity set to small (0.5), and the Resnik-normalized semantic similarity measure [44]. Differential protein expression analysis between RPMI-7951 si(-) and RPMI-7951 si(αV) datasets was performed using the QSpec spectral counter tool [45]. A volcano plot was generated in GraphPad Prism to visualize differentially expressed proteins, using a fold-change threshold of >1.5 (RPMI-7951 si(-) vs. RPMI-7951 si(αV)) and a -log(FDR) >1. Fold changes were calculated based on QSpec output values.

### Cell Survival Analysis

MTT (3-((4,5-dimethylthiazol-2-yl)-2,5-diphenyltetrazolium bromide) (Millipore) assay was used to determine sensitivity of cells to paclitaxel (PTX, Sigma-Aldrich). Briefly, 24 hours after seeding in 96-well tissue culture plates (1–1.2 × 10^4^ cells/well) cells were treated with different concentrations of PTX. Seventy-two hours later, the absorbance of MTT-formazan product dissolved in DMSO was measured with a microplate reader (Awareness Technology, Inc.) at 600 nm. Absorbance is proportional to the number of viable cells.

### Cell Migration

Serum-starved (24 hours) cells (8 × 10^4^) were placed in Transwell Cell Culture Inserts (pore size, 8 µm) (Corning) in DMEM containing 0.1% (w/v) BSA) and left to migrate toward 10% (v/v) FBS in DMEM as a chemoattractant. After 22 hours, cells that remained on the upper side of the inserts were removed with cotton-tipped swabs. Inserts were fixed in 4% paraformaldehyde for 15 minutes followed by staining with 1% (w/v) crystal violet in PBS for 90 minutes. Cells on the underside of the inserts were photographed using Olympus BX51TF microscope (five images/sample). The number of cells was determined using ImageJ software (Multi-point tool).

### Cell proliferation assay

Cell proliferation was analysed using the Click-iT assay according to the manufacturer’s instructions (Thermo Fisher Scientific, United States). Cells were seeded in a 6-well plate (4.5 × 10^5^ cells/well) and upon 24 h transfected with either control siRNA or specific siRNA. After 24 h, cells were trypsinised and seeded back in the same well to mimic steps during MTT assay. Next day, two hours before harvesting, modified thymidine analogue EdU (5-ethynyl-2’-deoxyuridine, final concentration 10 µM) was added. Cells were collected, fixed with 4% (w/v) paraformaldehyde, permeabilized with saponin, stained with Alexa-Fluor 488 azide (in the presence of CuSO4) and analysed by flow cytometry. Flow cytometry was performed using BD FACSCalibur (BD Biosciences, United States). Data were analysed using FCS Express (De Novo Software, United States). To determine the proliferation rate, the frequencies of the proliferative (EdU +) cells were compared.

### Confocal Microscopy

For immunofluorescence analysis (IF), cells were plated on coverslips in a 24-well plate (3.7 × 10^4^ cells/well). After 48 hours, cells were fixed using ice-cold methanol for 10 minutes or 2% (v/v) paraformaldehyde for 12 minutes followed by permeabilization with 0.1% (v/v) Triton X-100 for 2 minutes, and incubated with specific primary antibodies for 1 hour, followed by conjugated secondary antibodies for 1 hour (See Supplementary Table S1, Additional File 1). Actin stress fibers were stained with rhodamine phalloidin (Cell Signaling Technology). Cells were mounted with DAPI Fluoromount-G (SouthernBiotech). Fluorescence and respective Interference Reflection Microscopy (IRM) images were acquired using an inverted confocal microscope Leica TCS SP8 X (Leica Microsystems) with the HC PL APOCS2 63×/1.40 oil-immersion objective, zoom set at 2×. Images were analysed using LAS X software 3.1.1 (Leica Microsystems) and ImageJ (NIH, USA). For actin stress fibers quantification, only fibers that ended with FAs, marked by FA marker staining (analysed but not presented in figures), were identified as stress fibers and quantified. For MT quantification, only MTs within 5 µm from the cell edge (identified through IRM images) were quantified. All images were taken with the focus adjusted to the adhesion sites of cells at the upper surface of glass coverslip.

### Proximity ligation assay

Proximity ligation assay (PLA), using Duolink® PLA technology [46], was done according to manufacturer’s instructions. Briefly, cells seeded on coverslips for 48 hours were fixed, blocked and incubated with selected primary antibodies for 1 hour at +37 °C in a preheated humidity chamber (See Supplementary Table S1, Additional File 1). Following washing, coverslips are incubated in secondary antibodies conjugated to proprietary oligonucleotide arms (Navenibodies) for 1 hour. Incubation with specific enzymes enables the formation and amplification of a DNA circle in the spots of protein proximity. Fluorescent dots, generated by binding of fluorescently labelled probes, were visualized using an inverted confocal microscope (Leica TCS SP8 X, Leica Microsystems) with the HC PL APOCS2 63×/1.40 oil-immersion objective, zoom set at 2×. Images were analysed using LAS X software 3.1.1 (Leica Microsystems) and ImageJ (NIH, USA).

### Live Cell Imaging

For time-lapse live cell microscopy, RPMI-7951-EB3 cells were plated in 4-chamber 35 mm Cellvis glass bottom dishes (4.5 × 10^4^ cells/chamber). Twenty-four hours later, cells were transfected with either control siRNA or specific siRNA and imaged 48 hours after transfection. Images were taken every 33 seconds for five min per cell using HC PL APO CS2 63×/1.40 oil-immersion objective on the Leica TCS SP8 X microscope (Leica Microsystems), 3.5× zoom. The obtained movies were analysed with ImageJ software (Manual tracking plugin).

### Statistical Analysis

GraphPad Prism version 9.0.0 (GraphPad Software) was used to analyse data. Data obtained from IF, PLA and migration assay were analysed by one-way analysis of variance (ANOVA) with Dunnett’s multiple comparison or by unpaired Welch’s t-test, and expressed as histograms depicting mean ± standard deviation (SD); violin plots or scatter plots with marked median. Data obtained from MTT experiments were analysed by two-way ANOVA with Šídák’s multiple comparisons test, with a single pooled variance and expressed as mean ± SD. ns denotes not significant; * denotes p < 0.05; ** denotes p < 0.01; *** denotes p < 0.001; **** denotes p < 0.0001.

## RESULTS

### Melanoma cell line RPMI-7951 primarily utilizes integrin heterodimer ***α***V***β***5 for adhesion

IACs, together with secreted ECM proteins, were isolated from RPMI-7951 melanoma cells grown for 72 hours as described previously [28, 33]. The optimal crosslinking duration was determined based on intracellular IAC components (liprin β1, FAK and paxillin) (See Supplementary Fig. S1, S7, Additional File 2). MS analysis identified 624 proteins, of which 45 are components of the consensus adhesome and 388 of the meta adhesome [47, 48] (See Supplementary Table S2.1, Additional File 3).

The adhesome of RPMI-7951 cells contained three integrin receptor subunits: αV, β5 and β1. The most abundant integrin subunits were αV and β5, suggesting that these cells primarily use αVβ5 and possibly αVβ1 for adhesion in long-term 2D culture. α5 was detected, but at very low levels, suggesting that these cells may also express α5β1 (See Supplementary Table S2.1, Additional File 3).

PPI network of the identified proteins indicated that the majority of identified proteins were ECM proteins and actin-binding proteins (Fig. 1A; See Supplementary Table S2.2, Additional File 3). Proteins reported to bind MTs, intermediate filament-related proteins, enzymes, HSP proteins and nuclear proteins were also identified. GO enrichment analysis was performed using DAVID [41, 42], and proteins were assigned to functional groups and visualized using REVIGO [44]. Analysis indicated a significant enrichment of GO terms related to ECM, extracellular exosome and FA (Fig. 1B).

**Fig. 1.**
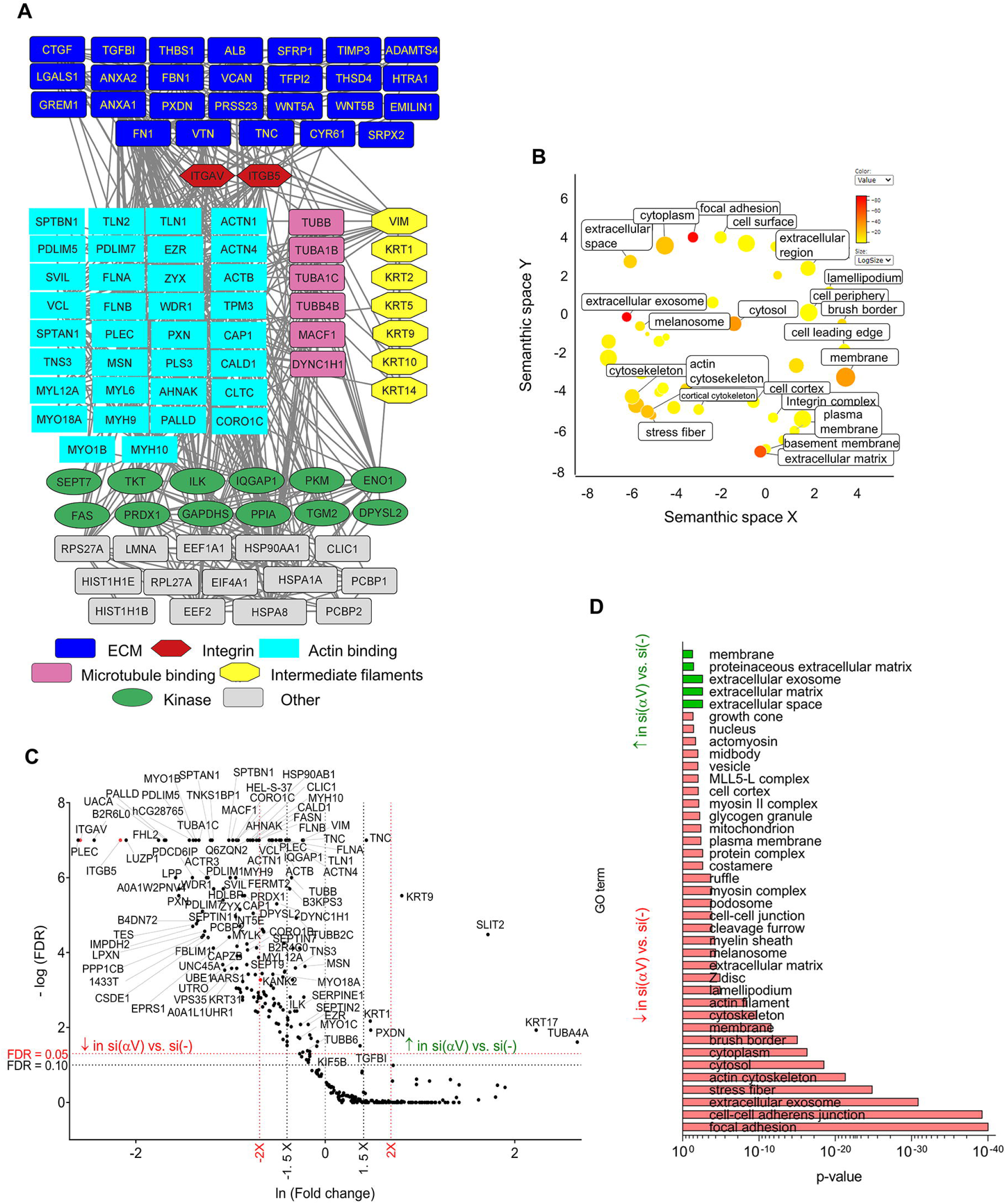
Mass spectrometry analysis of IACs isolated from RPMI-7951 cells. **(A)** Protein–protein interaction network of proteins identified by MS in IACs isolated from RPMI-7951 cells. Shapes represent identified proteins and are labelled with gene symbols, arranged and coloured according to their functional group as indicated. **(B)** IACs isolated from RPMI-7951 are enriched with proteins connected to the ECM, extracellular exosome and FA. Proteins from (A) (number of spectral counts ≥4, FDR < 5%, probability for protein identification ≥99.9%) were annotated using the DAVID GO database and visualized using the REVIGO tool, where p-values related to GO terms of cellular components (GOTERM_CC_DIRECT), represented by the colour bar and size of the circle. Statistically significant GO terms (p > 0.05) are presented. **(C)** Volcano plot analysis of proteins detected in IACs isolated from RPMI-7951 cells transiently transfected with non-specific siRNA (si(-)) versus integrin αV-specific siRNA (si(αV)). IAC proteins are visualized as volcano plot after the analysis with QSpec/QProt. To determine the significantly changed proteins −Log (FDR) ≥ 1 (red horizontal dotted line) corresponding to FDR ≤ 0.05 and −Log (FDR) ≥ 1,3 (black horizontal dotted line) corresponding to FDR ≤ 0.1; and fold change ≥ 1.5 (black vertical dotted line) or 2 (red vertical dotted line) were used. Upper left quadrant – proteins detected with lower levels of spectra, upper right quadrant – proteins detected with higher levels of spectra. Only proteins identified with a minimal number of spectral counts ≥ 4 in at least one biological replicate, FDR < 5%, probability for protein identification ≥ 99.9% were visualized. **(D)** DAVID GO analysis of significant proteins from (C). Statistically significant GO terms were presented in reverse x-axis of Benjamini corrected p-value (the smaller the p-value - the bigger the length of the bar), with higher (green arrow) and lower (red arrow) abundances in si(αV) then si(-).↓, decreased abundance, ↑, increased abundance.

### Integrin αVβ5-induced interaction networks

To determine which proteins were found in αVβ5-containing adhesions, transient knockdown of integrin subunit αV was performed and IAC composition was compared to control siRNA treatment using QSpec/QProt [45]. Differences were visualized using a volcano plot (Fig. 1C). Integrin αV knockdown removed virtually all αVβ5 from the cell surface, but did not alter β1 levels (See Supplementary Table S2.3, Additional File 3), indicating that RPMI-7951 cells use other alpha subunit(s), possibly α5, to pair with β1. A volcano plot, displaying proteins whose abundance was different in RPMI-7951 cells transfected with integrin αV-specific siRNA compared to control siRNA (Fig. 1C), identified key components of integrin αVβ5 IACs dependent on αV. The expression of 164 proteins was decreased by αV knockdown (See Supplementary Table S2.3, Additional File 3).

Proteins that were elevated following integrin αV knockdown included slit homolog 2 protein (SLIT2), peroxidasin homolog (PXDN), tenascin C (TNC) and transforming growth factor-β-induced protein ig-h3 (TGFBI). DAVID analysis connected most of the proteins with extracellular space, ECM, extracellular exosome, proteinaceous extracellular matrix and membrane (Fig. 1D, See Supplementary Table S2.3, Additional File 3).

Proteins detected at lower levels upon integrin αV knockdown and detected by high number of spectra (See Supplementary Table S2.3, Additional File 3) were actin-binding adhesome proteins (plectin, vinculin, talins 1 and 2, α-actinins 1 and 4, filamin A and B, PDLIM7 and PDLIM5), non-muscle myosins 9 and 10 (MYH9 and MYH10), zyxin (ZYX), ezrin (EZR), moesin (MSN), tensin 3 (TNS3), tropomyosin 3 isoform 1 (TPM3), palladin (PALLD) and caldesmon 1 (CALD1). Actin (ACTB), which was identified with a very high number of spectra, was also greatly reduced upon integrin αV knockdown. Furthermore, the levels of Ras GTPase-activating-like protein 1 (IQGAP1) and integrin linked kinase (ILK), which have been identified in consensus adhesome, were reduced [47]. DAVID analysis connected most of the proteins with FAs, cell-cell adherens junctions, extracellular exosome, stress fibers and actin cytoskeleton (Fig. 1D, See Supplementary Table S2.3, Additional File 3). Selected MS results were confirmed by WB and/or IF (for plectin, talin1, talin2, vinculin, filamin A and B, α-actinins 1 and 4, IQGAP1) (See Supplementary Fig. S2A, S8, S2C, Additional File 2). It has to be noted that in the adhesome we found TNS3. TNS3, together with TNS1 and TNS2, are found in FAs and FBs in fibroblasts, although to different degrees. TNS2 is localized mainly in FAs, while TNS3 is mostly found in FBs, and TNS1 is found in both [49]. Our data, which show the presence of integrin α5 and β1, albeit in a small number of spectra, and TNS3, support the possibility that RPMI-7951 cells, under long-term growth conditions, may contain FBs.

Finally, a reduced abundance of a few proteins that are not recognized as adhesome proteins was found. These included the chloride intracellular channel 1 (CLIC1), which was recently identified to recruit PIP5K1A/C to the leading edge of the plasma membrane, to generate a PIP2-rich microdomain and to activate talin [50].

### Differential role of KANK2 in PTX sensitivity and migration in two melanoma cell lines, MDA-MB-435S and RPMI-7951

The CMSC proteins MACF1, liprin β1 and KANK2 were detected in RPMI-7951 cell IACs (See Supplementary Table S2.2, Additional File 3), and were reduced upon transient integrin αV knockdown (See Supplementary Table S2.3, Additional File 3). Interestingly, no KANK1 was detected. Other CMSC proteins, liprin α1 and ELKS were represented at a lower abundance, while CLASPs (CLIP-associating proteins), kinesin family member 21A (KIF21A), plus-end tracking protein EB1 and LL5β were not found in RPMI-7951 cells, either transfected with control or integrin αV siRNA (See Supplementary Table S2.3, Additional File 3). The reduced abundance of liprin β1 and KANK2 was confirmed in RPMI-7951 cells transfected with integrin αV-specific siRNA using WB (See Supplementary Fig. S2B, S9, Additional File 2).

Since we previously demonstrated that RPMI-7951 cells, similar to MDA-MB-435S cells, can be sensitized to microtubule poison PTX through integrin αV or β5 knockdown [51], we investigated whether talin2 and KANK2 similarly influence actin-microtubule crosstalk and thereby regulate PTX sensitivity [28, 33]. Before addressing this, we confirmed that talin2 or KANK2 knockdown does not affect cell proliferation (Fig. 2A) which is similar to results previously obtained in MDA-MB-435S cells [28]. Talin2 knockdown in RPMI-7951 cells increased PTX sensitivity, as indicated by the greater survival difference between control and talin2 knockdown cells in the presence of PTX (Fig. 2B), mimicking integrin αV or β5 knockdown [51]. Surprisingly, transient KANK2 knockdown in RPMI-7951 cells had no effect on PTX sensitivity (Fig. 2C). This differs from the effect observed in MDA-MB-435S cells, where talin2 and KANK2 functionally interact within integrin αvβ5 FAs and knockdown of either protein sensitized the cells to microtubule poison PTX [28]. The differential roles of KANK2 between these cell models was also evident in cell migration. In MDA-MB-435S cells, knockdown of either talin2 or KANK2 reduced cell migration, thus mimicking the effect of integrin αV or β5 knockdown [28]. However, in RPMI-7951 cells, talin2 knockdown mimicked the effect of integrin αV or β5 knockdown and reduced cell migration, while KANK2 knockdown enhanced cell migration (Fig. 2D). We conclude that KANK2 plays distinct roles in regulating PTX sensitivity and cell migration between two melanoma cell lines, MDA-MB-435S and RPMI-7951. Therefore, we decided to further investigate KANK2-containing adhesions in the RPMI-7951 cell line.

**Fig. 2.**
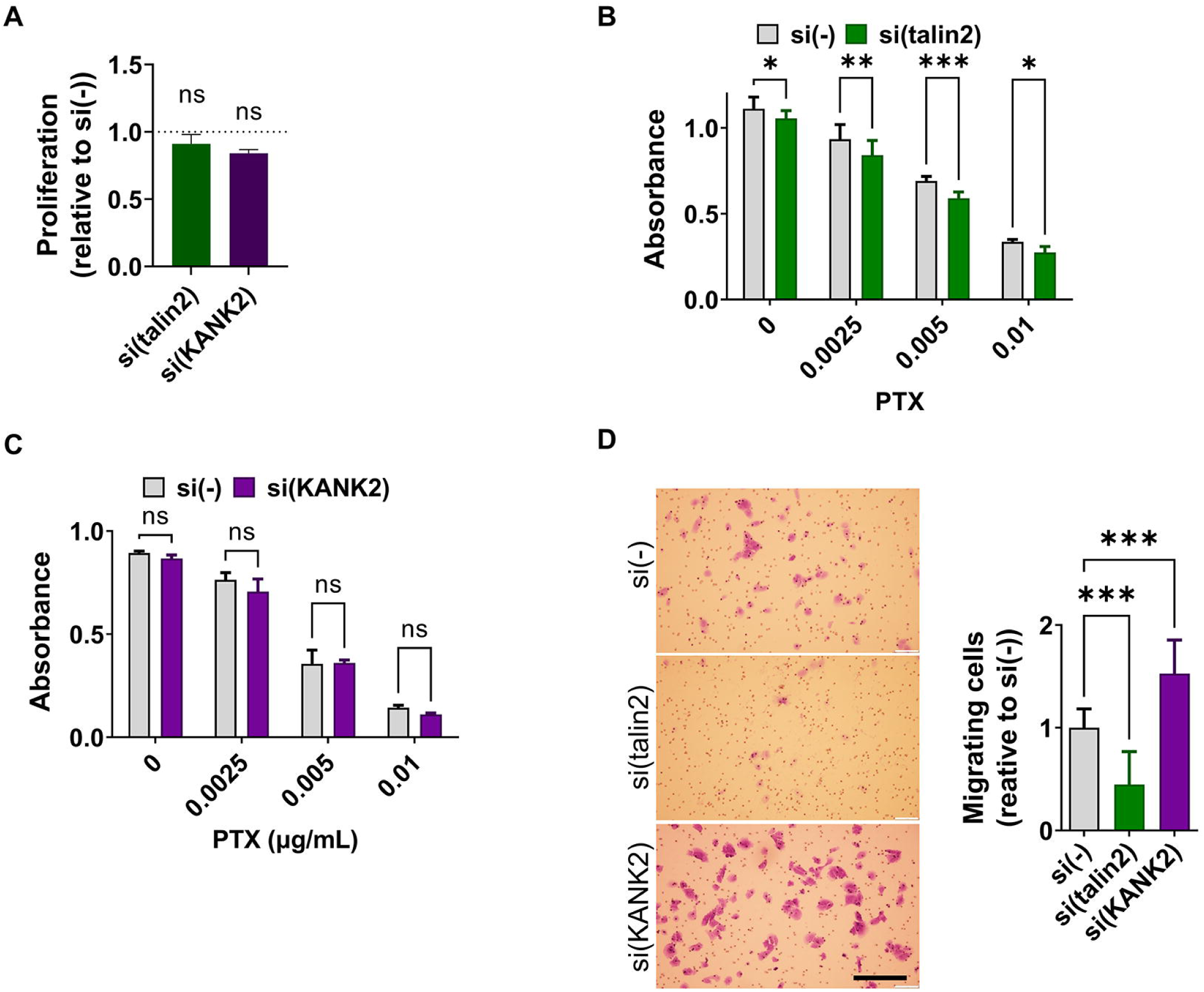
Talin2 and KANK2 differentially affect cell sensitivity to PTX and cell migration. (**A**) Neither talin2 nor KANK2 knockdown influences cell proliferation. Cell proliferation was measured using ClickIT EdU assay upon transfection with either control, talin2 or KANK2-specific siRNA. Histogram represents measurements of percentage of EdU + cells of si(talin2) or si(KANK2) relative to si(-), plotted as mean ± SD (n = 2, > 1000 cells per experiment). Data were analyzed by one-way ANOVA with Dunnett’s multiple comparison. ns, not significant; *P < 0.05; **P < 0.01; ***P < 0.001; ****P<0.0001. In RPMI-7951 cells **(B)** talin2 knockdown increases sensitivity to PTX, while **(C)** KANK2 knockdown decreases it. Twenty-four hours upon transfection, cells were seeded in 96-well plates and 24 hours later treated with different concentrations of PTX. Cytotoxicity was measured by MTT assay. Data were analysed by two-way analysis of variance (ANOVA) with Šídák’s multiple comparisons test, with a single pooled variance. ns, not significant; *P < 0.05; **P < 0.01; ***P < 0.001; ****P < 0.0001. (n = 3). **(D)** Talin2 knockdown decreases, and KANK2 knockdown increases migration in RPMI-7951 cells. Serum starved (24 hours) cells, transfected previously with either control or talin2-specific siRNA were seeded in Transwell cell culture inserts and left to migrate for 22 hours toward serum. Cells on the underside of the inserts were stained with crystal violet, photographed, and counted. Scale bar = 100 μm. Histogram data represents averages of five microscope fields of three independently performed experiments, plotted as mean ± SD. Data were analysed by unpaired Student’s t-test. ns, not significant; *P < 0.05; **P < 0.01; ***P < 0.001; ****P < 0.0001.

### In RPMI-7951 cells talin1 is required for maintenance of integrin ***α***V***β***5 FAs while talin2 regulates FAs dynamics

The MS-proteomic data above demonstrate that integrin αV knockdown leads to the loss of talin1, talin2 and KANK2. To determine the localization of talin isoforms within integrin αVβ5 IACs, IF was employed. Figure 3A shows that both talin1 and talin2 colocalize with integrin β5. Talin1 colocalizes with integrin β5 at the cell edge in structures that resemble FAs. Integrin β5 colocalizes with talin2 both at the cell edge in FAs and in the center of the cell in structures which are talin1-negative, and according to localization and appearance represent RAs [4, 33]. Proximity of integrin β5 and talin2 was further confirmed using PLA (Fig. 3B). This is in line with our MS data that both talins are part of integrin αVβ5 IACs.

**Fig. 3.**
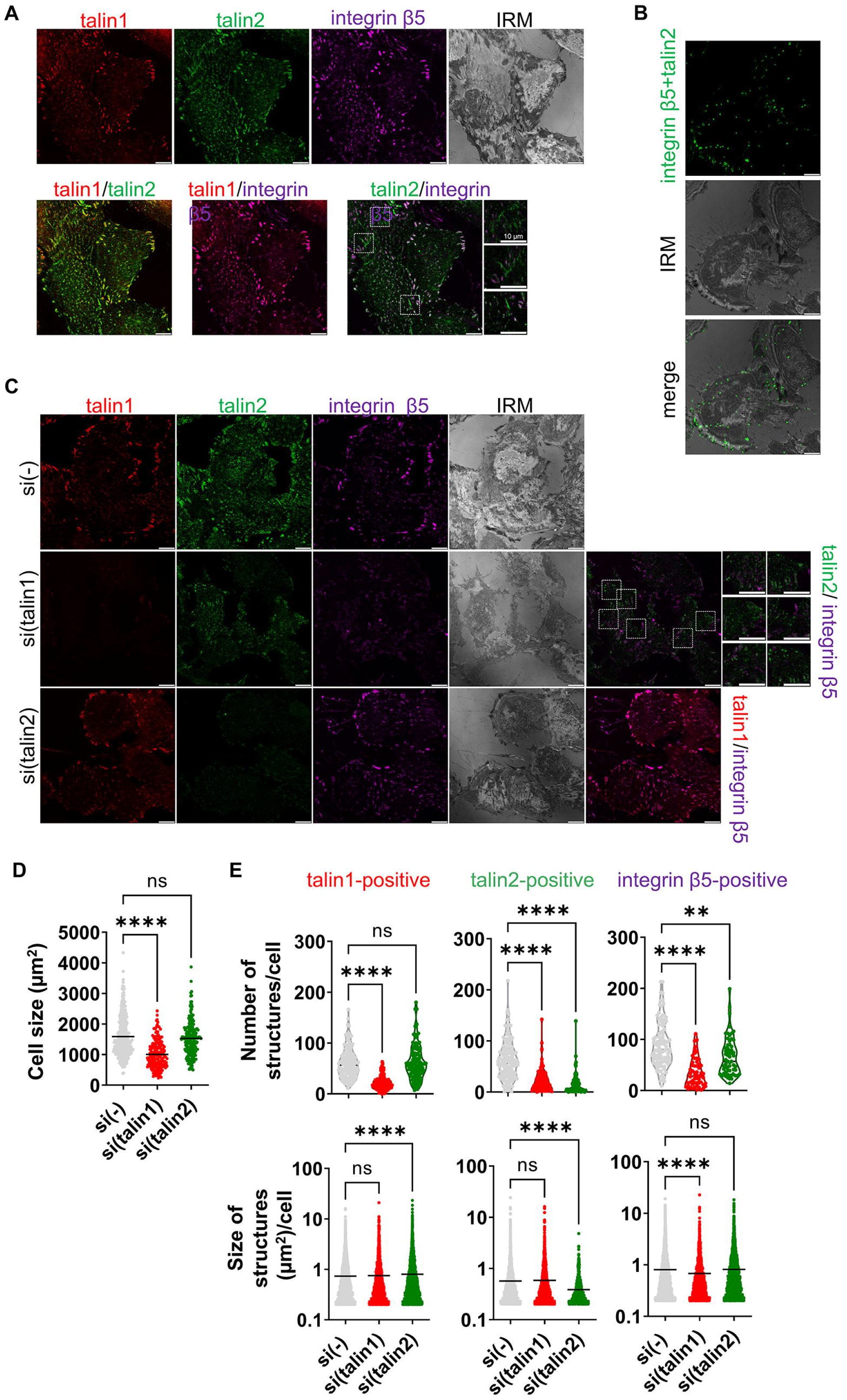
Talin1 is necessary for maintenance of integrin αVβ5 FAs, while talin2 influences their dynamics. **(A)** Talin1 and talin2 are localized within integrin αVβ5 FAs. Forty-eight hours after seeding, RPMI-7951 cells were methanol fixed, and stained with anti-talin1 antibody followed by Alexa-Fluor IgG1 555-conjugated antibody (red), anti-talin2 antibody followed by Alexa-Fluor IgG2b 488-conjugated antibody (green) and anti-integrin β5 antibody followed by Alexa-Fluor 647-conjugated antibody (magenta) and IRM images were taken. Analysis was performed using TCS SP8 Leica. Scale bar = 10 μm. **(B)** Integrin β5 and talin2 are in close proximity. Forty-eight hours after seeding RPMI-7951 cells were methanol fixed, and a PLA assay with anti-integrin β5 and anti-talin2 primary antibody was performed. Generated fluorescent dots were visualized and IRM images were taken. Analysis was performed using TCS SP8 Leica. Scale bar = 10 μm. **(C)** Talin1 knockdown, disrupts αVβ5 FAs and a reduces cell size, while talin2 knockdown only increased size of talin1-positive structures and reduced the number of integrin β5-positive structures. Forty-eight hours after transfection with either control siRNA, talin1 or talin2-specific siRNA, RPMI-7951 cells were methanol fixed, and stained with anti-talin1 antibody followed by Alexa-Fluor IgG1 555-conjugated antibody (red), anti-talin2 antibody followed by Alexa-Fluor IgG2b 488-conjugated antibody (green) and anti-β5 antibody followed by Alexa-Fluor 647-conjugated antibody (magenta) and IRM images were taken. Analysis was performed using TCS SP8 Leica. Scale bar = 10 μm. **(D, E)** Quantification of data presented in (C) Violin plots (number of structures/cell) and scatter plots with median marked in size (cell size, size of structures/cell) represents measurements of > 45 cells. Data were analysed by one-way ANOVA with Dunnett’s multiple comparison. ns, not significant; * P < 0.05; ** P < 0.01; *** P < 0.001; **** P < 0.0001.

It has been shown that talin2 also localizes to FBs [52]. Although RPMI-7951 cells were not seeded on the α5β1 ligand fibronectin, the MS analysis showed that cells excrete their own fibronectin and contain a low number of spectra specific for integrin heterodimers α5 and β1 in IACs isolates (See Supplementary Table S2.1, Additional File 3). Talin2 was found in rod-shaped αVβ5-negative structures (Fig. 3A, second row magnified images), which are likely FBs. In addition, in RPMI-7951 cells, the presence of α5-positive IACs that did not overlap with vinculin or integrin β5 was confirmed, indicating a structure different from FAs, i.e. FBs (See Supplementary Fig. S3, Additional File 2).

Next, RPMI-7951 cells were transfected with control, talin1- or talin2-specific siRNA and the location of both talins and integrin β5 was analysed. Knockdown of talin1 led to disruption of αVβ5 FAs and a reduction in cell size (Fig. 3C), supporting a key role for talin1 in maintaining FAs, as previously described by others [26]. Upon talin1 knockdown, a simultaneous reduction of talin2-positive structures was observed. The remaining talin2 signal, which still colocalized with integrin β5, was found in the center of the cells and likely represents RAs (Fig. 3C, middle row), which are reported to be more stable and contain talin2 [4, 16]. In addition, after talin1 knockdown, talin2-positive and β5-negative rod-shaped structures reminiscent of FBs were observed (Fig. 3C, middle row, magnified images). Atherton et al. [53] reported that the formation of FBs depends on force, but their maintenance is largely independent of intracellular tension. Therefore, it is possible that FBs remain upon talin1 knockdown. This conclusion was confirmed by analysing vinculin/talin1/talin2 expression in cells after talin1 knockdown where only RAs and FBs remain (See Supplementary Fig. S4, Additional File 2). After talin2 knockdown, talin1-positive and integrin β5-positive FAs were still present, mostly at the cell edge and there was no visible effect on cell size (Fig. 3C, last row).

A quantitative assessment following talin1 knockdown, which notably reduced cell size (Fig. 3D), demonstrated a significant reduction of number of talin2- or integrin β5-positive structures (Fig. 3E, first row), indicating the significant loss of integrin αVβ5 FAs. Simultaneously, the size of talin2-positive structures did not change (FBs and RAs) but the size of integrin β5-positive structures was greatly reduced (RAs) (Fig. 3E, second row). As stated above, talin2 knockdown did not affect cell size (Fig. 3D) nor the number of talin1-positive structures, however, their size increased (Fig. 3E), which indicates altered FAs dynamics. These FAs are unlikely to be αVβ5 FAs, as talin2 knockdown did not change the size integrin β5-positive structures (Fig. 3E), suggesting that the enlarged structures may be integrin α5β1 FAs, whose presence was indicated by MS analysis. Additionally, talin2 knockdown reduced the number of integrin β5-positive structures without changing their size, which could indicate the reduction of number of structures other than integrin αVβ5 FAs, likely RAs (Fig. 3E). Indeed, analysis of the levels of RA components Numb and AP2 in IAC samples following talin2 knockdown and cytochalasin D treatment revealed their reduced levels [54].

### In RPMI-7951 cells talin1 localizes with KANK2 predominantly at the cell edge while talin2 and KANK2 localize predominantly in the cell center

A key question was which talin isoform colocalizes with KANK2 in RPMI-7951 cells. Therefore, IF and PLA analysis of talin1, talin2 and KANK2 was performed. KANK2 was present in talin1- and talin2-positive structures at the cell edge (FAs) (Fig. 4A). The analysis of KANK2 proximity to talin1 also showed that they are predominantly located at the cell edge (Fig. 4B, C). KANK2 was also found in the central part of the cell where it colocalized mostly with talin2 but not with talin1, thus indicating structures different from FAs (Fig. 4A). Talin2-KANK2 proximity analysis showed the majority of positive signals located in the cell center (Fig. 4B, C).

**Fig. 4.**
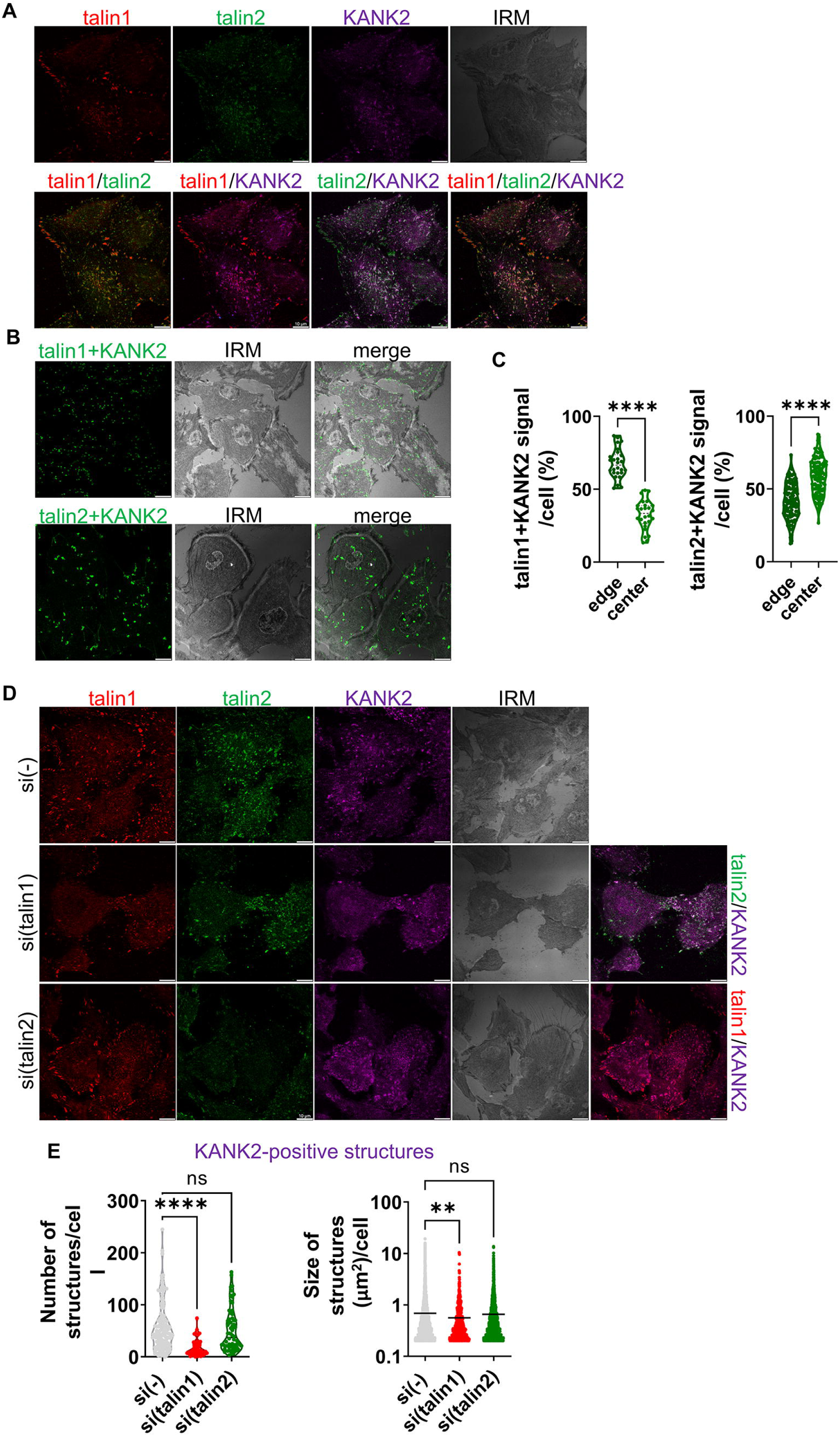
KANK2 localizes with talin1 at the cell edge, and with talin2 in the cell centre. **(A)** KANK2 localizes in talin1 and talin2-positive adhesions. Forty-eight hours after seeding RPMI-7951 cells were methanol fixed, and stained with anti-talin1 antibody followed by Alexa-Fluor IgG1 555-conjugated antibody (red), anti-talin2 antibody followed by Alexa-Fluor IgG2b 488-conjugated antibody (green) and anti-KANK2 antibody followed by Alexa-Fluor 647-conjugated antibody (magenta) and IRM images were taken. Analysis was performed using TCS SP8 Leica. Scale bar = 10 μm. **(B)** KANK2 is in close proximity to talin1 and talin2. Forty-eight hours after seeding cells were methanol fixed, and a PLA assay using anti-KANK2 combined with either anti-talin1 or anti-talin2 primary antibody performed. Generated fluorescent dots were visualized and IRM images were taken. Analysis was performed using TCS SP8 Leica. Scale bar = 10 μm. **(C)** Quantification of data presented in (B). Violin plots (percentage of positive signal/cell) represents measurements of > 30 cells. Data were analysed by one-way ANOVA with Dunnett’s multiple comparison. ns, not significant; * P < 0.05; ** P < 0.01; *** P < 0.001; **** P < 0.0001. **(D)** Knockdown of talin1, unlike talin2, leads to changes in KANK2 localization. Forty-eight hours after transfection with either control siRNA, talin1 or talin2-specific siRNA, RPMI-7951 cells were methanol fixed and stained with anti-talin1 antibody followed by Alexa-Fluor IgG1 555-conjugated antibody (red), anti-talin2 antibody followed by Alexa-Fluor IgG2b 488-conjugated antibody (green), anti-KANK2 antibody followed by Alexa-Fluor 647-conjugated antibody (magenta) and IRM images were taken. Analysis was performed using TCS SP8 Leica. Scale bar = 10 μm. **(E)** Quantification of data presented in (D). Violin plot (number of structures/cell) and scatter plot with marked median (cell size, size of structures/cell) represents measurements of > 45 cells. Data were analysed by one-way ANOVA with Dunnett’s multiple comparison. ns, not significant; * P < 0.05; ** P < 0.01; *** P < 0.001; **** P < 0.0001.

RPMI-7951 cells were then transfected with control, talin1- or talin2-specific siRNA and the location of both talins and KANK2 was analysed. Knockdown of talin1 reduced the number and size of KANK2-positive structures (Fig. 4D, E). Since the knockdown of talin1 disrupts FAs and reduces the number of talin2-positive structures (Fig. 3C, E), the remaining KANK2-structures which colocalize with talin2 (Fig. 4D) are either Ras [54] or FBs [53].

As before (Fig. 3C, E), the talin2 knockdown did not affect the number of talin1-positive structures but enhanced their size, indicating alteration of FA dynamics. The knockdown of talin2 does not exhibit the same effect on KANK2-positive structures, as it does not alter their number or size (Fig. 4D, E). This does not imply that the knockdown of talin2 has no effect on KANK2 function. In fact, we previously demonstrated in MDA-MB-435S cells that disruption of talin2-KANK2 interaction upon talin2 knockdown disconnects FAs from CMSCs but this disruption does not impact the localization of KANK2 [28].

### In RPMI-7951 cells KANK2 is part of all three adhesions: FAs, RAs and FBs

As previously emphasized, KANK2 is localized in talin1-positive adhesions at the cell edge, but the majority of KANK2 is localized in the central region of the cell, where it colocalizes only with talin2 (Fig. 4). We hypothesized that these central structures are α5β1 FBs and αVβ5 RAs. Our new data identified KANK2 within αVβ5 RAs [54]. Next, to determine whether KANK2 localizes to FBs, the colocalization of integrin α5 and KANK2 was analysed. The results demonstrated that they indeed predominantly colocalize in the central part of the cell within rod-shaped structures i.e. FBs (Fig. 5A). The proximity between integrin α5 and KANK2 was further confirmed through PLA (Fig. 5B). Therefore, in addition to being in FAs and RAs, KANK2 is also part of FBs.

**Fig. 5.**
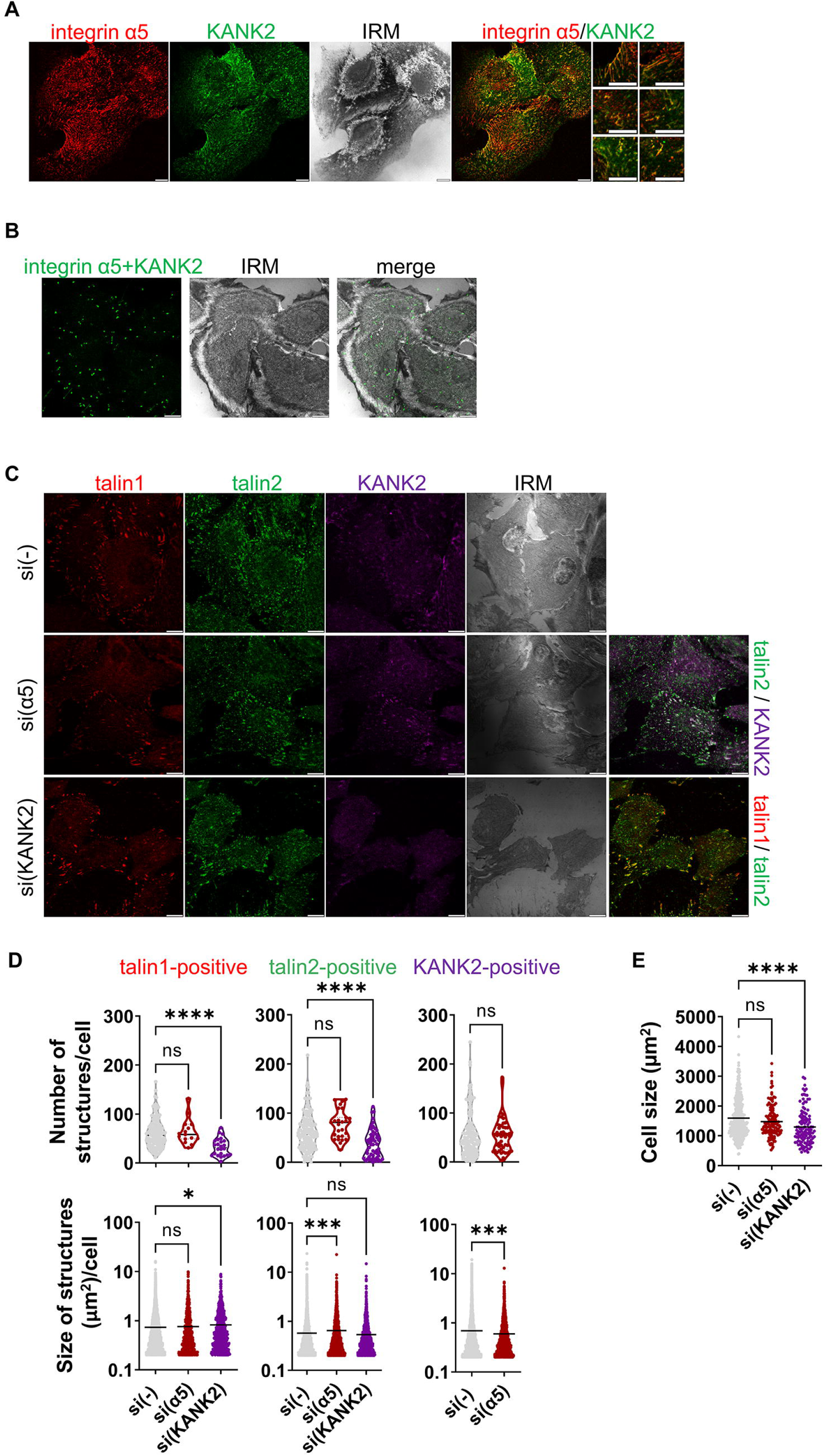
In RPMI-7951 cells KANK2 is part of FAs, FBs, RAs. **(A)** KANK2 localizes in integrin α5-positive structures. Forty-eight hours after seeding RPMI-7951 cells were fixed and permeabilised, and stained with anti-integrin α5 antibody followed by Alexa-Fluor 546-conjugated antibody (red), anti-KANK2 antibody followed by Alexa-Fluor 488-conjugated antibody (green) and IRM images were taken. Analysis was performed using TCS SP8 Leica. Scale bar = 10 μm. **(B)** KANK2 is in close proximity with integrin α5. Forty-eight hours after seeding cells were PFA fixed and permeabilised, and a PLA assay using anti-KANK2 combined with anti-integrin α5 primary antibody was performed. Generated fluorescent dots were visualized and IRM images were taken. Analysis was performed using TCS SP8 Leica. Scale bar = 10 μm. **(C)** Effect of knockdown of integrin α5 or KANK2 on cell size, the number and size of talin1 and talin2 structures. Forty-eight hours after transfection with either control siRNA, integrin α5 or KANK2-specific siRNA, RPMI-7951 cells were methanol fixed and stained with anti-talin1 antibody followed by Alexa-Fluor IgG1 555-conjugated antibody (red), anti-talin2 antibody followed by Alexa-Fluor IgG2b 488-conjugated antibody (green), anti-KANK2 antibody followed by Alexa-Fluor 647-conjugated antibody (magenta) and IRM images were taken. Analysis was performed using TCS SP8 Leica. Scale bar = 10 μm. **(D, E)** Quantification of data presented in (C). Violin plots (number of structures/cell) and scatter plots with median marked in size (cell size, size of structures/cell) represents measurements of > 45 cells. Data were analysed by one-way ANOVA with Dunnett’s multiple comparison. ns, not significant; * P < 0.05; ** P < 0.01; *** P < 0.001; **** P < 0.0001.

To analyse how a reduction of FBs affects other IACs, RPMI-7951 cells were transfected with control-or integrin α5-specific siRNA and the localization of talin1, talin2 and KANK2 was analysed. The knockdown of integrin α5 did not affect cell size (Fig. 5C, E), the number of talin1-, talin2-, or KANK2-positive structures, or the size of talin1-positive structures (FAs) (Fig. 5C, D). However, the size of talin2-positive structures increased, while size of KANK2-positive structures decreased (Fig. 5C, D), indicating a change in the dynamics of talin2-and KANK2-positive adhesions. The fact that FAs, FBs and RAs contain talin2 and KANK2 suggest that there is a crosstalk between FBs, FAs and/or RAs.

### In RPMI-7951 cells KANK2-talin1 functional interaction is required for FA maintenance while KANK2-talin2 functional interaction regulates FA dynamics

Published studies [4, 28, 53], our new data [54] combined with our own findings presented here (Fig. 3A-C; Fig. 4A, B; Fig. 5A, B) demonstrate that KANK2 and talin2 are part of FAs, RAs and FBs. Since we have shown above that talin1 regulates FA maintenance while talin2 affects FA dynamics, we aimed to determine the role of KANK2 in these events.

The knockdown of KANK2 reduced cell size (Fig. 5C, E), and the number of talin1-positive structures, but increased their size indicating change in the dynamics of FAs formed by either integrins αVβ5 or α5β1 (Fig. 5C, D). KANK2 knockdown also increased the size of α5-positive, but not β5-positive, structures (Fig. 6A-D), implying altered dynamics of α5β1 FA and/or FBs. However, this does not rule out the possibility that KANK2 knockdown influences αVβ5-positive adhesion dynamics, as αVβ5 forms FA and RAs which are very different in size, therefore overall changes in size may not be easily detected.

**Fig. 6.**
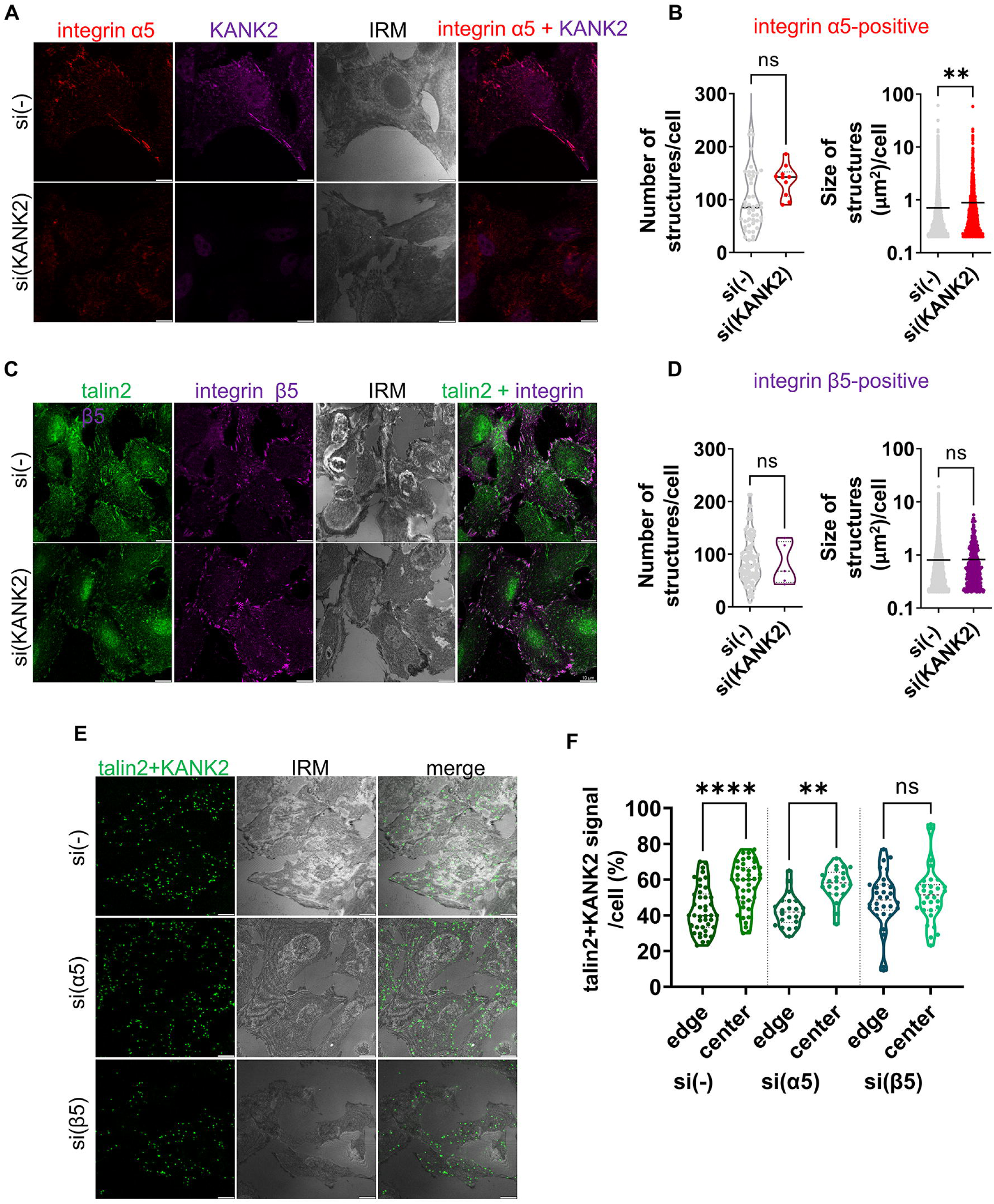
In RPMI-7951 cells KANK2 is in close proximity to talin2 in FAs, FBs, RAs. **(A-D)** KANK2 knockdown increases integrin α5-positive structures, but has no effect on integrin β5-positive structures. Forty-eight hours after transfection with either control siRNA or KANK2-specific siRNA RPMI-7951 cells were fixed and permeabilised, and stained with (A) anti-integrin α5 antibody followed by Alexa-Fluor 546-conjugated antibody (red) and anti-KANK2 antibody followed by Alexa-Fluor 488-conjugated antibody (green) or (C) anti-talin2 antibody followed by Alexa-Fluor IgG2b 488-conjugated antibody (green) and anti-KANK2 antibody followed by Alexa-Fluor 647-conjugated antibody (magenta). IRM images were taken. Analysis was performed using TCS SP8 Leica. Scale bar = 10 μm. (B, D) Quantification of data presented in (A, C respectively). Violin plots (number of structures/cell) and scatter plots with median marked in size (cell size, size of structures/cell) represents measurements of > 45 cells. Data were analysed by one-way ANOVA with Dunnett’s multiple comparison. ns, not significant; * P < 0.05; ** P < 0.01; *** P < 0.001; **** P < 0.0001. **(E, F)** Knockdown of integrin α5 or integrin β5 influences the distribution of talin2 KANK2 proximity signal in RPMI-7951 cells. (E) Forty-eight hours after transfection with either control siRNA, integrin α5- or integrin β5-specific siRNA RPMI-7951 cells were methanol fixed and a PLA assay using anti-KANK2 combined with anti-talin2 primary antibody was performed. Generated fluorescent dots were visualized and IRM images were taken. Analysis was performed using TCS SP8 Leica. Scale bar = 10 μm. (F) Quantification of data presented in (E). Violin plots (percentage of positive signal/cell) represents measurements of > 20 cells. Data were analysed by one-way ANOVA with Dunnett’s multiple comparison. ns, not significant; * P < 0.05; ** P < 0.01; *** P < 0.001; **** P < 0.0001.

The reduction in the number of FAs after KANK2 knockdown (according to decreased number of talin1-positive structures) aligns with the observed reduction in cell size (Fig. 5C, E), which was previously also noted after talin1 knockdown (Fig. 3D). Since talin1 knockdown reduces talin2-positive structures (Fig. 3C, E), it was expected that KANK2 knockdown, which reduces talin1-positive structures, would also reduce the number of talin2-positive structures; indeed, this was the case (Fig. 5C, D). In conclusion, KANK2 knockdown mimics the effect of talin1 knockdown which indicates that the functional interaction between KANK2 and talin1 is necessary for FA maintenance. The knockdown of KANK2 mimics the effect of talin2 knockdown on the size of talin1-positive structures (increased size), suggesting that KANK2-talin2 functional interaction affects FA dynamics. Since talin2 knockdown only increases the size of talin1-positive structures (Fig. 3C, E) and doesn’t affect either number or size of α5- (data not shown) or β5-positive structures (Fig. 3C, E), we cannot conclude in which FAs KANK2-talin2 functionally interact.

### Talin2-KANK2 are in proximity in FAs, RAs and FBs

Next, PLA was employed to analyse talin2-KANK2 colocalization following integrin α5 or β5 knockdown. We hypothesized that we would observe a difference in distribution (cell edge vs cell center) as a consequence of loss of either FAs and RAs (integrin β5 knockdown) or integrin α5 FAs and FBs (integrin α5 knockdown). The distribution (percentage of talin2-KANK2-positive PLA signals at the cell edge or cell center) in cells transfected with control siRNA showed preferential location of talin2-KANK2-positive PLA signals in the cell center. After integrin α5 knockdown a small reduction in talin2-KANK2-positive PLA signals was observed in the cell center and a slight increase at the cell edge (Fig. 6E, F). This is in line with a reduction of FBs. After β5 knockdown, the percentage of talin2-KANK2-positive PLA signals at the cell edge and cell center changed even more, i.e. a strong reduction of signal in the cell center and increase at the cell edge were observed (Fig. 6E, F). This is in line with a reduction of αVβ5 FA and RA, leaving only α5-positive FAs and FBs. In summary, talin2-KANK2 are in proximity to each other in FAs, RAs and FBs.

### Loss of integrin ***α***5 mimics the effect of KANK2 and increases cell migration

Since we found that KANK2 is located in FBs and that KANK2 knockdown increases migration, we hypothesized that α5 knockdown could mimic the effect of KANK2. Indeed, integrin α5 knockdown increased cell migration (Fig. 7A, B). To support the conclusion that increased migration is the consequence of reduced KANK2 from FBs and not αVβ5 FAs or RAs, migration was also analysed after β5 knockdown and showed that it had the opposite effect, i.e. reduction of migration (See Supplementary Fig. S5A, B, Additional file 2). Migration depends on the cytoskeleton, especially actin and MTs [55]. Therefore, actin and MT distribution was analysed after talin1, talin2, integrin α5 or KANK2 knockdown using IF (Fig. 7C). The only change in actin was following talin1 knockdown which disrupted actin stress fibers (Fig. 7C, D); however, analysis of MTs showed that KANK2 and integrin α5 knockdown led to a more compact appearance of the MT network at the cell edge (Fig. 7C, E). Talin2 knockdown, which reduces cell migration (Fig. 2C), led to a lower amount of MTs at the cell edge, which is the opposite effect of KANK2 or α5 knockdown (Fig. 7C, E). The observed changes in MT organization, as well as the changes in adhesion size following talin2 or KANK2 knockdown (Fig. 3E; 5C, D), indicate altered MT dynamics. Therefore, we isolated an RPMI-7951 cell population transfected with fluorescent EB3 (RPMI-7951-EB3), which enabled us to follow growing MTs during time-lapse live cell microscopy, since fluorescent EB3 (EB3-Dendra2) binds to the growing tip of MTs [56]. By measuring the velocity of MT growth upon talin2, integrin α5 and KANK2 knockdown using live cell imaging of fluorescent EB3, we showed that the velocity of MT plus end growth upon α5 knockdown was approximately 2 times faster than in EB3 cells transfected with control siRNA (Fig. 7F; See Supplementary Fig. S6, Additional File 2; Supplementary Movie S1, Additional file 4; Supplementary Movie S3, Additional file 6). While knockdown of talin2 and KANK2 had the same effect as α5 knockdown, the difference in the velocity of MT growth was more pronounced upon KANK2 knockdown than upon talin2 knockdown (1.46 and 1.23 times faster than the control, respectively) (Fig. 7F; See Supplementary Fig. S6, Additional File 2; Supplementary Movie S1, Additional file 4; Supplementary Movie S4, Additional file 7; Supplementary Movie S2, Additional file 5). Since integrin α5 knockdown decreased the size of KANK2-positive structures (Fig. 5C, D), we hypothesize that this loss of KANK2 results in fewer MTs in the cell center and more MTs at the cell edge. Because integrin α5 knockdown did not affect the number or size of talin1-positive structures (FAs) (Fig. 5D), it is more likely that the enhanced motility is due to the loss of FBs rather than changes in FAs dynamics. Nevertheless, our results clearly show that KANK2 is a component of FBs, acting as a link to MTs and contributing to their stabilisation. The loss of integrin α5 or KANK2 from FBs, which increases cell migration, involves the action of MT cytoskeleton.

**Fig. 7.**
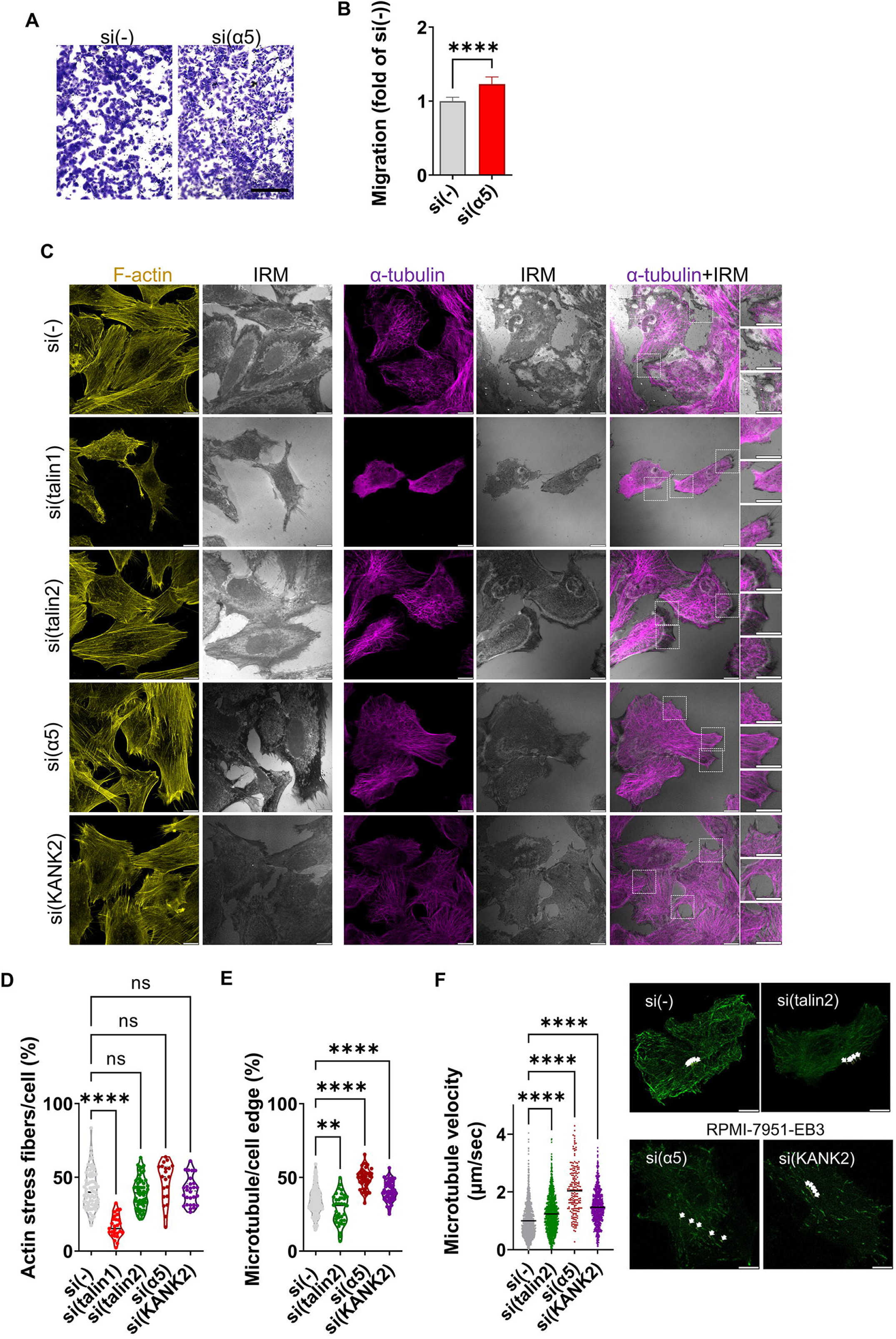
Loss of integrin α5 or KANK2 from FBs increases cell migration through changes in microtubules. **(A)** Integrin α5 knockdown increases migration in RPMI-7951 cells. Serum starved (24 hours) cells, transfected previously with control or integrin α5-specific siRNA, were seeded in Transwell cell culture inserts and left to migrate for 22 hours toward serum. Cells on the insert underside were stained with crystal violet, photographed, and counted. Scale bar = 100 μm. **(B)** Histogram data represents averages of five microscope fields of three independently performed experiments, plotted as mean ± SD. Data were analysed by unpaired Student’s t-test. ns, not significant; *P < 0.05; **P < 0.01; ***P < 0.001; ****P < 0.0001. **(C)** Upon KANK2 or integrin α5 knockdown MTs are denser at the cell edge. Forty-eight hours after transfection with control, talin1, talin2, integrin α5 or KANK2-specific siRNA. Cells were fixed with methanol (α-tubulin visualization) or PFA followed by permeabilization with Triton X-100 (F-actin visualization), stained with anti-α-tubulin antibody followed by Alexa-Flour 647-conjugated antibody (magenta) or incubated with Alexa-Flour 488 conjugated phalloidin (shown in gold) and IRM images were taken. Analysis was performed using TCS SP8 Leica. Scale bar = 10 μm. **(D, E)** Quantification of data from (C). Violin plots represents measurements of > 40 cells. Data were analysed by unpaired Student’s t-test. ns, not significant; *P < 0.05; **P < 0.01; ***P < 0.001; ****P < 0.0001. **(F)** MT growth velocity increased upon talin2, integrin α5, or KANK2 knockdown. Violin plot represents quantification of time-lapse live cell microscopy measurements of > 450 analysed microtubules, (n = 3) relative to velocity of MT growth in cells transfected with control siRNA (set as 1). Data were analysed by one-way ANOVA with Dunnett’s multiple comparison test. ns, not significant; *P < 0.05; **P < 0.01; ***P < 0.001; ****P < 0.0001. Still images represent tracking of one microtubule tip in 132 s through five frames.

## DISCUSSION

Metastasis remains the primary cause of mortality in melanoma patients [57]. Current treatment options, including targeted drug therapy and immunotherapy, are not universally effective, with some patients exhibiting resistance. Therefore, better understanding of metastatic melanoma is required for the development of effective treatments. Integrins αVβ5 and α5β1 have been identified as potential therapeutic targets [5, 57–59]. However, clinical trials targeting integrins have thus far been unsuccessful [57, 60, 61] due to several possible reasons: (i) variable integrin expression, (ii) redundancy in integrin function, (iii) distinct roles of integrins at various disease stages, and (iv) sequestering of therapeutics by integrin-containing tumor-derived extracellular vesicles [5–7]. We hypothesize that targeting IACs, which act downstream of integrins, may provide a more consistent therapeutic outcome.

In two melanoma cell lines, MDA-MB-435S and RPMI-7951, we showed that integrin αV or β5 knockdown increases sensitivity to the MT poison PTX and reduces cell migration [51, this work]. To better understand the composition of IACs and their connection to observed changes in cell migration and PTX sensitivity, we analysed the adhesome of both cell lines as well as the adhesome following integrin αV knockdown in each line. Analysis of the MDA-MB-435S adhesome revealed that integrin αV knockdown reduces the abundance of its downstream effectors, KANK2 and talin2 [33]. These proteins functionally interact, and their individual knockdown mimics integrin αV or β5 knockdown, increasing PTX sensitivity and reducing cell migration [28].

Here we analysed the adhesome of RPMI-7951 cells and found that, in long-term culture, these cells also preferentially use integrin αVβ5 for adhesion. However, we also found integrin subunits α5 and β1, which form α5β1 FAs and FBs, and confirmed their presence by IF. Integrin αV knockdown in RPMI-7951 cells reduced KANK2 and talin2 levels but only talin2 mimicked the effect of integrin αV or β5 knockdown on PTX sensitivity and migration. In contrast, KANK2 knockdown increased cell migration. This led us to further investigate KANK2 localization and function in RPMI-7951 cell adhesions.

Another difference between MDA-MB-435S and RPMI-7951 cells was the impact of integrin αV knockdown in the expression of ECM proteins, specifically TNC and TGFβ1. In MDA-MB-435S cells, integrin αV knockdown slightly reduced the abundance of these proteins [33]. In RPMI-7951 cells, however, integrin αV knockdown increased the abundance of both proteins. These data suggest a differential feedback loop where cells secrete more of the ECM components necessary to ensure a homeostatic adhesion when some key adhesion molecules are missing. Given that TNC and TGFβ1 are linked to melanoma invasion and metastasis [62, 63], this differential response highlights the complexity of integrin-mediated regulation in melanoma progression.

In RPMI-7951 cells, talin1 and talin2 colocalize with integrin β5 at the cell edge within FAs. Talin2 is also found in rod-shaped αVβ5-negative structures (FBs) and αVβ5-positive structures in the cell center (RAs). Similar patterns were observed in MDA-MB-435S cells [28], with the key exception that these cells, under growth conditions on uncoated substrates, form much less, if any FBs [54].

The reasons for the different localization of talin1 and talin2 in cells remain unknown. Both talins bind to β-integrin cytoplasmic tails *via* their N-terminal FERM domains, but the talin2 has a slightly higher binding affinity than talin1 [64, 65]. In RPMI-7951 cells, talin1 is essential for FA formation, as its knockdown leads to FA disassembly, and a subsequent loss of talin2. In contrast, talin2 knockdown does not affect FA number (talin1-positive adhesions) but reduces the number of integrin β5-positive adhesions, later identified as RAs [54]. While talin2 is a component of RAs, it is not essential for their formation [4, 54]. Integrin αVβ5 and talin2 found in RAs can also be recruited to FAs, a process enhanced by increased cellular tension [19]. Since in RPMI-7951 cells talin2 knockdown increases the size of talin1-positive adhesions without affecting actin stress fibers, it is unclear whether the reduction of RAs is due to talin2 exiting RAs or from changes in cell tension, as indicated by FA increased size.

As noted above, talin2 knockdown increases the size of talin1-positive adhesions, indicating changes in adhesion dynamics. However, no increase in size of β5- or α5-positive adhesions was observed. We assume that knockdown of talin2 affects primarily β5-positive adhesions. This is partly because statistical analysis of β5-positive adhesion size, observed through IF, may be unreliable, as it includes both FAs and RAs.

In RPMI-7951, as in MDA-MB-435S cells [28], there was no compensation between talin1 and talin2. A similar lack of compensation between talins was observed in early embryonic development [66, 67]. However, in fibroblasts, talin2 can compensate for the loss of talin1 [68], and in endothelial cells, which express only talin1, introduced talin2 can compensate for talin1 loss [69].

In HeLa cells and the HaCaT keratinocyte cell line, talin1 binds to KANK1, triggering the formation of CMSC, which surrounds adhesion sites and regulates their formation and dynamics via MT-dependent signalling and trafficking [25]. In fibroblasts, KANK2 localizes to both FAs and in central sliding adhesions, where it interacts with talin1, dynein light chain isoforms (Dynll1 and Dynll2) and liprin-β1 [27]. Recently, the proximity of TNS3 (enriched in FBs) to both talin1 and 2 and KANK2 was demonstrated [53]. In MDA-MB-435S cells, KANK2 functionally interacts with talin2 within integrin αVβ5 FAs, playing a role in actin-MT dynamics [28].

Our results in RPMI-7951 cells suggests a link between KANK2 and talin1 in FAs. KANK2 knockdown mimics talin1 knockdown by reducing cell size. It also decreased the number of talin1-positive adhesions (FAs). Since talin1 is a main regulator of FA assembly, this reduction subsequently leads to decreased talin2 levels and enlarged remaining FAs. PLA analysis revealed that KANK2-talin1 positive signals are predominantly located at the cell edge, where FAs are positioned. In contrast, KANK2-talin2 signals are located predominantly in the cell center, corresponding to the location of FBs and RAs. We tried to obtain additional evidence of which talin binds KANK2 using WB analysis of isolated IACs after talin1 or 2 knockdowns, as was previously done in MDA-MB-435S cells where talin2 knockdown significantly reduced cross-linked KANK2 levels, confirming its binding to talin2 [28]. However, in RPMI-7951 cells, cross-linked KANK2 levels after talin2 knockdown remained inconclusive (data not shown). The results obtained in MDA-MB-435S cells [28] were informative as these cells form FAs and RAs and do not contain FBs [54]. However, in RPMI-7951 cells talin2 and KANK2 are present in all three types of adhesions, FAs, FBs and RAs, thus adding complexity to the analysis.

In RPMI-7951 FBs, KANK2 is linked to talin2, as demonstrated by talin2-KANK2 PLA analysis following integrin α5 or β5 knockdown. Upon integrin α5 knockdown, the percentage of talin2-KANK2 PLA signals in the cell center decreases, consistent with a reduction in FBs. Following β5 knockdown, the percentage of talin2-KANK2 PLA signals at the cell edge and cell center changes compared to control, leaving only α5-positive FAs and FBs. Since the precise distribution of talin2-KANK2 levels in FAs and RAs remain unknown, the observed changes in the cell center and at the cell edge indicate that talin2 and KANK2 are present in both types of β5-positive adhesions. Collectively, our findings indicate that in RPMI-7951 cells, KANK2 associates with talin1 in FAs and with talin2 in all three adhesion types - FAs, RAs and FBs. However, KANK2 shows a clear preference for talin1 in FAs, whereas in FBs and RAs, which lack talin1, KANK2 is linked to talin2.

The factors determining why KANK2 interacts with talin1 or talin2 are not known. One possibility is that KANK proteins generally localize to the periphery of FAs [25, 27]. This suggests that FAs may initially form with talin1, while talin2 is later recruited to the outer rim, allowing KANK2 from CMSC to establish actin-MT crosstalk [70]. In MDA-MB-435S cells, we hypothesized that talin1 is phosphorylated by CDK1 [71] and therefore has a low KANK2 binding affinity. It is unclear whether the situation is reversed in RPMI-7951 cells, where talin2 might be phosphorylated, leading KANK2 to favour talin1.

In both MDA-MB-435S and RPMI-7951 cells, integrin αV or β5 knockdown increases PTX sensitivity and reduces cell migration [51]. However, while KANK2 and talin2 knockdown in MDA-MB-435S cells mimic this effect [28, 33], we found that KANK2 knockdown in RPMI-7951 cells has a different outcome. It does not affect PTX sensitivity and unexpectedly increases cell migration. This finding aligns with studies in fibroblasts, where adhesion sliding and formation of FBs containing KANK2 was linked to reduced migration speed [27]. It also supports our observation that KANK2 knockdown mimics the integrin α5 knockdown, leading to increased migration.

In RPMI-7951 cells KANK2 knockdown increased the size of α5- and talin1-positive structures indicating altered α5β1 FA and/or FBs dynamics. It also mimicked integrin α5 knockdown by increasing cell migration and altering MTs, specifically increasing the percentage of MTs at the cell edge. This contrasts with findings in MDA-MB-435S cells, where talin2 or KANK2 knockdown from αVβ5 FAs disrupted actin-MT crosstalk, leading to decreased MT presence at the cell edge [28]. Talin2 knockdown in RPMI-7951 cells, despite colocalizing with KANK2 and integrin α5 FBs, decreased the percentage of MTs at the cell edge and mimicked integrin αV or β5 knockdown by reducing cell migration. It has been shown that MT destabilization promotes the relocalization of integrin β5 from flat clathrin lattices, which are interspersed within RAs [72], to FAs through tension modulation [19]. Therefore, our study highlights the fact that in RPMI-7951 cells talin2, KANK2 and integrin α5 knockdown influence MTs. Confirmation of altered MT dynamics was obtained by observing an increase in velocity of MT growth. It remains unclear whether these effects are a result of direct connection between FBs and MTs (e.g. through CMSCs) or crosstalk between different adhesion structures. Further investigation is needed to clarify how FBs interact with MTs.

After integrin α5 knockdown, talin2-positive structures increased in size, while size of KANK2-positive structures decreased. Since talin1- and β5-positive structures remained unaffected, it is difficult to hypothesize which adhesions were impacted. However, since talin2 and KANK2 are present in all three adhesion structures FA, FB and RA, we conclude there is crosstalk between them. Indeed, it has been recently demonstrated that RAs are opposed by active integrin α5β1 [73].

In conclusion, the functional interactions between talin1, talin2 and KANK2 appear to be cell type-specific. However, our findings reveal that these interactions are influenced by their location within a specific adhesion structure. The factors guiding talin1/talin2 and KANK2 to particular adhesions remain unknown. To unlock the potential of integrins to be targets in metastatic melanoma, particularly for limiting metastasis, it is important to understand IAC composition and the role of individual components in different adhesions. Our study shows that KANK2 is present in distinct adhesion structures and likely has different binding partners. Future studies should focus on identifying interactors both *in vitro* and *in vivo* to uncover new avenues for cancer treatment.

## List of abbreviations

ECM: Extracellular matrix
IAC: Integrin adhesion complex
FA: Focal adhesion
FB: Fibrillar adhesion
RA: Reticular adhesion
MT: Microtubule
CMSC: Cortical microtubule stabilizing complex
KANK: KN motif and ankyrin repeat domain containing protein
ABS2 F-: actin binding site 2
DMEM: Dulbecco’s Modified Eagle Medium
FBS: Fetal Bovine Serum
ITGAV: Integrin Subunit αV
ITGA5: Integrin Subunit α5
ITGB5: Integrin Subunit β5
HEPES: Hydroxyethylpiperazine ethanesulfonic acid
DTBP: Dimethyl 3,3’-dithiobispropionimidate, Wang and Richard’s reagent
MS: Mass spectrometry
STRING: Search Tool for the Retrieval of Interacting Genes/Proteins
MTT 3-: ((4,5-Dimethylthiazol-2-yl)-2,5-diphenyltetrazolium bromide
PTX: Paclitaxel
DMSO: Dimethyl Sulfoxide
PBS: Phosphate-Buffered Saline
IRM: Interference reflection microscopy
IF: Immunofluorescence analysis
PLA: Proximity ligation assay
WB: Western blotting
SD: Standard Deviation
ANOVA: One-way analysis of variance
RhoA GEF ARHGEF2 / GEF-H1: RhoA Guanine Nucleotide Exchange Factor
PPI: network Protein–Protein Interaction Network
HSP: proteins Heat Shock Proteins
GO: Gene Ontology
DAVID: Database for Annotation, Visualization and Integrated Discovery
REVIGO: Reduce and Visualize Gene Ontology
SLIT2: Slit Homolog 2 Protein
PXDN: Peroxidasin Homolog
TNC: Tenascin C
TGFBI: Transforming Growth Factor-β-Induced Protein IG-H3
PDLIM7: PDZ and LIM Domain Protein 7
PDLIM5: PDZ and LIM Domain Protein 5
MYH9/MYH10: Non-Muscle Myosin Heavy Chain 9 / 10
ZYX: Zyxin
EZR: Ezrin
MSN: Moesin
TNS3: Tensin 3
TPM3: Tropomyosin 3, Isoform 1
PALLD: Palladin
CALD1: Caldesmon 1
ACTB: Beta-Actin
IQGAP1: Ras GTPase-Activating-Like Protein 1
ILK: Integrin-Linked Kinase
CLIC1: Chloride Intracellular Channel 1
MACF1: Microtubule-Actin Cross-Linking Factor 1
ELKS: Protein Rich in Glutamate (E), Leucine (L), Lysine (K), and Serine (S)
CLASP: CLIP-Associated Protein
KIF21A: Kinesin Family Member 21A
LL5β PHLDB2: (Pleckstrin Homology Like Domain Family B Member 2), commonly referred to as LL5β

## Declarations

### Ethics approval and consent to participate

Not applicable

### Consent for publication

Not applicable

### Availability of data and materials

The datasets supporting the conclusions of this article are available in the ProteomeXchange Consortium via the PRIDE partner repository, dataset identifier PXD064756, https://www.ebi.ac.uk/pride/archive/. Other raw data supporting the conclusions of this article will be made available by the authors, without undue reservation, to any qualified researcher.

### Author contributions

Material preparation, data collection and analysis were performed by [NS], [AR], [ML], [AT], [MP], [MA], and [AA-R]. [NS] and [AR] contributed equally to the experimental work. The first draft of the manuscript was written by [NS] and [AA-R], corrections were made by [AR], [ML], [JDH] and [MJH]. All authors read and approved the final manuscript. [NS] and [AA-R] equally contributed to the study conception and design.

## Acknowledgements

We thank Marina Šutalo of the Laboratory for Cell Biology and Signalling for technical assistance.

## Authors’ information (optional)

Not applicable

## REFERENCES

1. Green HJ, Brown NH. Integrin intracellular machinery in action. Exp Cell Res. 2019;378(2):226–31.

2. Burridge K, Guilluy C. Focal adhesions, stress fibers and mechanical tension. Exp Cell Res. 2016;343(1):14–20.

3. Walko G, Castanon MJ, Wiche G. Molecular architecture and function of the hemidesmosome. Cell Tissue Res. 2015;360(2):363–78.

4. Lock JG, Jones MC, Askari JA, Gong X, Oddone A, Olofsson H, et al. Reticular adhesions are a distinct class of cell-matrix adhesions that mediate attachment during mitosis. Nat Cell Biol. 2018;20(11):1290–302.

5. Pang X, He X, Qiu Z, Zhang H, Xie R, Liu Z, et al. Targeting integrin pathways: mechanisms and advances in therapy. Signal Transduct Target Ther. 2023;8(1):1.

6. Bergonzini C, Kroese K, Zweemer AJM, Danen EHJ. Targeting Integrins for Cancer Therapy - Disappointments and Opportunities. Front Cell Dev Biol. 2022;10:863850.

7. Samarzija I, Dekanic A, Humphries JD, Paradzik M, Stojanovic N, Humphries MJ, et al. Integrin Crosstalk Contributes to the Complexity of Signalling and Unpredictable Cancer Cell Fates. Cancers (Basel). 2020;12(7).

8. Geiger B, Yamada KM. Molecular architecture and function of matrix adhesions. Cold Spring Harb Perspect Biol. 2011;3(5).

9. Winograd-Katz SE, Fassler R, Geiger B, Legate KR. The integrin adhesome: from genes and proteins to human disease. Nat Rev Mol Cell Biol. 2014;15(4):273–88.

10. Yamaguchi N, Knaut H. Focal adhesion-mediated cell anchoring and migration: from in vitro to in vivo. Development. 2022;149(10).

11. Mishra YG, Manavathi B. Focal adhesion dynamics in cellular function and disease. Cell Signal. 2021;85:110046.

12. Zamir E, Katz M, Posen Y, Erez N, Yamada KM, Katz BZ, et al. Dynamics and segregation of cell-matrix adhesions in cultured fibroblasts. Nat Cell Biol. 2000;2(4):191–6.

13. Katz BZ, Zamir E, Bershadsky A, Kam Z, Yamada KM, Geiger B. Physical state of the extracellular matrix regulates the structure and molecular composition of cell-matrix adhesions. Mol Biol Cell. 2000;11(3):1047–60.

14. Pankov R, Cukierman E, Katz BZ, Matsumoto K, Lin DC, Lin S, et al. Integrin dynamics and matrix assembly: tensin-dependent translocation of alpha(5)beta(1) integrins promotes early fibronectin fibrillogenesis. J Cell Biol. 2000;148(5):1075–90.

15. Conway JRW, Jacquemet G. Cell matrix adhesion in cell migration. Essays Biochem. 2019;63(5):535–51.

16. Zuidema A, Wang W, Kreft M, Te Molder L, Hoekman L, Bleijerveld OB, et al. Mechanisms of integrin alphaVbeta5 clustering in flat clathrin lattices. J Cell Sci. 2018;131(21).

17. Zuidema A, Wang W, Sonnenberg A. Crosstalk between Cell Adhesion Complexes in Regulation of Mechanotransduction. Bioessays. 2020;42(11):e2000119.

18. Baschieri F, Dayot S, Elkhatib N, Ly N, Capmany A, Schauer K, et al. Frustrated endocytosis controls contractility-independent mechanotransduction at clathrin-coated structures. Nat Commun. 2018;9(1):3825.

19. Zuidema A, Wang W, Kreft M, Bleijerveld OB, Hoekman L, Aretz J, et al. Molecular determinants of alphaVbeta5 localization in flat clathrin lattices - role of alphaVbeta5 in cell adhesion and proliferation. J Cell Sci. 2022;135(11).

20. Alfonzo-Mendez MA, Sochacki KA, Strub MP, Taraska JW. Dual clathrin and integrin signaling systems regulate growth factor receptor activation. Nat Commun. 2022;13(1):905.

21. Klapholz B, Brown NH. Talin - the master of integrin adhesions. J Cell Sci. 2017;130(15):2435–46.

22. Qi L, Jafari N, Li X, Chen Z, Li L, Hytonen VP, et al. Talin2-mediated traction force drives matrix degradation and cell invasion. J Cell Sci. 2016;129(19):3661–74.

23. del Rio A, Perez-Jimenez R, Liu R, Roca-Cusachs P, Fernandez JM, Sheetz MP. Stretching single talin rod molecules activates vinculin binding. Science. 2009;323(5914):638–41.

24. Goult BT, Brown NH, Schwartz MA. Talin in mechanotransduction and mechanomemory at a glance. J Cell Sci. 2021;134(20).

25. Bouchet BP, Gough RE, Ammon YC, van de Willige D, Post H, Jacquemet G, et al. Talin-KANK1 interaction controls the recruitment of cortical microtubule stabilizing complexes to focal adhesions. Elife. 2016;5.

26. Goult BT, Yan J, Schwartz MA. Talin as a mechanosensitive signaling hub. J Cell Biol. 2018;217(11):3776–84.

27. Sun Z, Tseng HY, Tan S, Senger F, Kurzawa L, Dedden D, et al. Kank2 activates talin, reduces force transduction across integrins and induces central adhesion formation. Nat Cell Biol. 2016;18(9):941–53.

28. Loncaric M, Stojanovic N, Rac-Justament A, Coopmans K, Majhen D, Humphries JD, et al. Talin2 and KANK2 functionally interact to regulate microtubule dynamics, paclitaxel sensitivity and cell migration in the MDA-MB-435S melanoma cell line. Cell Mol Biol Lett. 2023;28(1):56.

29. Chen NP, Sun Z, Fassler R. The Kank family proteins in adhesion dynamics. Curr Opin Cell Biol. 2018;54:130–6.

30. van der Vaart B, van Riel WE, Doodhi H, Kevenaar JT, Katrukha EA, Gumy L, et al. CFEOM1-associated kinesin KIF21A is a cortical microtubule growth inhibitor. Dev Cell. 2013;27(2):145–60.

31. Rafiq NBM, Nishimura Y, Plotnikov SV, Thiagarajan V, Zhang Z, Shi S, et al. Publisher Correction: A mechano-signalling network linking microtubules, myosin IIA filaments and integrin-based adhesions. Nat Mater. 2019;18(7):770.

32. Tadijan A, Samarzija I, Humphries JD, Humphries MJ, Ambriovic-Ristov A. KANK family proteins in cancer. Int J Biochem Cell Biol. 2021;131:105903.

33. Paradzik M, Humphries JD, Stojanovic N, Nestic D, Majhen D, Dekanic A, et al. KANK2 Links alphaVbeta5 Focal Adhesions to Microtubules and Regulates Sensitivity to Microtubule Poisons and Cell Migration. Front Cell Dev Biol. 2020;8:125.

34. Jones MC, Humphries JD, Byron A, Millon-Fremillon A, Robertson J, Paul NR, et al. Isolation of integrin-based adhesion complexes. Curr Protoc Cell Biol. 2015;66:9 8 1–9 8 15.

35. Robertson J, Jacquemet G, Byron A, Jones MC, Warwood S, Selley JN, et al. Defining the phospho-adhesome through the phosphoproteomic analysis of integrin signalling. Nat Commun. 2015;6:6265.

36. Nesvizhskii AI, Keller A, Kolker E, Aebersold R. A statistical model for identifying proteins by tandem mass spectrometry. Anal Chem. 2003;75(17):4646–58.

37. Deutsch EW, Bandeira N, Perez-Riverol Y, Sharma V, Carver JJ, Mendoza L, et al. The ProteomeXchange consortium at 10 years: 2023 update. Nucleic Acids Res. 2023;51(D1):D1539–D48.

38. Perez-Riverol Y, Bandla C, Kundu DJ, Kamatchinathan S, Bai J, Hewapathirana S, et al. The PRIDE database at 20 years: 2025 update. Nucleic Acids Res. 2025;53(D1):D543–D53.

39. Doncheva NT, Morris JH, Gorodkin J, Jensen LJ. Cytoscape StringApp: Network Analysis and Visualization of Proteomics Data. J Proteome Res. 2019;18(2):623–32.

40. Shannon P, Markiel A, Ozier O, Baliga NS, Wang JT, Ramage D, et al. Cytoscape: a software environment for integrated models of biomolecular interaction networks. Genome Res. 2003;13(11):2498–504.

41. Huang da W, Sherman BT, Lempicki RA. Bioinformatics enrichment tools: paths toward the comprehensive functional analysis of large gene lists. Nucleic Acids Res. 2009;37(1):1–13.

42. Huang da W, Sherman BT, Lempicki RA. Systematic and integrative analysis of large gene lists using DAVID bioinformatics resources. Nat Protoc. 2009;4(1):44–57.

43. Thomas PD, Campbell MJ, Kejariwal A, Mi H, Karlak B, Daverman R, et al. PANTHER: a library of protein families and subfamilies indexed by function. Genome Res. 2003;13(9):2129–41.

44. Supek F, Bosnjak M, Skunca N, Smuc T. REVIGO summarizes and visualizes long lists of gene ontology terms. PLoS One. 2011;6(7):e21800.

45. Choi H, Fermin D, Nesvizhskii AI. Significance analysis of spectral count data in label-free shotgun proteomics. Mol Cell Proteomics. 2008;7(12):2373–85.

46. Alam MS. Proximity Ligation Assay (PLA). Curr Protoc Immunol. 2018;123(1):e58.

47. Horton ER, Byron A, Askari JA, Ng DHJ, Millon-Fremillon A, Robertson J, et al. Definition of a consensus integrin adhesome and its dynamics during adhesion complex assembly and disassembly. Nature cell biology. 2015;17(12):1577–87.

48. Horton ER, Humphries JD, James J, Jones MC, Askari JA, Humphries MJ. The integrin adhesome network at a glance. J Cell Sci. 2016;129(22):4159–63.

49. Clark K, Howe JD, Pullar CE, Green JA, Artym VV, Yamada KM, et al. Tensin 2 modulates cell contractility in 3D collagen gels through the RhoGAP DLC1. J Cell Biochem. 2010;109(4):808–17.

50. Peng JM, Lin SH, Yu MC, Hsieh SY. CLIC1 recruits PIP5K1A/C to induce cell-matrix adhesions for tumor metastasis. J Clin Invest. 2021;131(1).

51. Stojanovic N, Dekanic A, Paradzik M, Majhen D, Ferencak K, Ruscic J, et al. Differential Effects of Integrin alphav Knockdown and Cilengitide on Sensitization of Triple-Negative Breast Cancer and Melanoma Cells to Microtubule Poisons. Mol Pharmacol. 2018;94(6):1334–51.

52. Praekelt U, Kopp PM, Rehm K, Linder S, Bate N, Patel B, et al. New isoform-specific monoclonal antibodies reveal different sub-cellular localisations for talin1 and talin2. Eur J Cell Biol. 2012;91(3):180–91.

53. Atherton P, Konstantinou R, Neo SP, Wang E, Balloi E, Ptushkina M, et al. Tensin3 interaction with talin drives the formation of fibronectin-associated fibrillar adhesions. J Cell Biol. 2022;221(10).

54. Rac A, Loncaric M, Stojanovic N, Fatima M, Resetar M, Hrsak D, et al. Regulation of reticular adhesions by KANK2 and talin2 in two melanoma cell lines. bioRxiv. 2025:2025.06.02.657402.

55. Schmidt CJ, Stehbens SJ. Microtubule control of migration: Coordination in confinement. Curr Opin Cell Biol. 2024;86:102289.

56. Stepanova T, Slemmer J, Hoogenraad CC, Lansbergen G, Dortland B, De Zeeuw CI, et al. Visualization of microtubule growth in cultured neurons via the use of EB3-GFP (end-binding protein 3-green fluorescent protein). J Neurosci. 2003;23(7):2655–64.

57. Arias-Mejias SM, Warda KY, Quattrocchi E, Alonso-Quinones H, Sominidi-Damodaran S, Meves A. The role of integrins in melanoma: a review. Int J Dermatol. 2020;59(5):525–34.

58. Vogetseder A, Thies S, Ingold B, Roth P, Weller M, Schraml P, et al. alphav-Integrin isoform expression in primary human tumors and brain metastases. Int J Cancer. 2013;133(10):2362–71.

59. Doyle AD, Nazari SS, Yamada KM. Cell-extracellular matrix dynamics. Phys Biol. 2022;19(2).

60. O’Day S, Pavlick A, Loquai C, Lawson D, Gutzmer R, Richards J, et al. A randomised, phase II study of intetumumab, an anti-alphav-integrin mAb, alone and with dacarbazine in stage IV melanoma. Br J Cancer. 2011;105(3):346–52.

61. Ruffini F, Graziani G, Levati L, Tentori L, D’Atri S, Lacal PM. Cilengitide downmodulates invasiveness and vasculogenic mimicry of neuropilin 1 expressing melanoma cells through the inhibition of alphavbeta5 integrin. Int J Cancer. 2015;136(6):E545–58.

62. Nummela P, Lammi J, Soikkeli J, Saksela O, Laakkonen P, Hölttä E. Transforming Growth Factor Beta-Induced (TGFBI) Is an Anti-Adhesive Protein Regulating the Invasive Growth of Melanoma Cells. The American Journal of Pathology. 2012;180(4):1663–74.

63. Shao H, Kirkwood JM, Wells A. Tenascin-C Signaling in melanoma. Cell Adh Migr. 2015;9(1-2):125–30.

64. Anthis NJ, Wegener KL, Ye F, Kim C, Goult BT, Lowe ED, et al. The structure of an integrin/talin complex reveals the basis of inside-out signal transduction. EMBO J. 2009;28(22):3623–32.

65. Anthis NJ, Wegener KL, Critchley DR, Campbell ID. Structural diversity in integrin/talin interactions. Structure. 2010;18(12):1654–66.

66. Monkley SJ, Zhou XH, Kinston SJ, Giblett SM, Hemmings L, Priddle H, et al. Disruption of the talin gene arrests mouse development at the gastrulation stage. Dev Dyn. 2000;219(4):560–74.

67. Monkley SJ, Kostourou V, Spence L, Petrich B, Coleman S, Ginsberg MH, et al. Endothelial cell talin1 is essential for embryonic angiogenesis. Developmental Biology. 2011;349(2):494–502.

68. Zhang X, Jiang G, Cai Y, Monkley SJ, Critchley DR, Sheetz MP. Talin depletion reveals independence of initial cell spreading from integrin activation and traction. Nat Cell Biol. 2008;10(9):1062–8.

69. Kopp PM, Bate N, Hansen TM, Brindle NPJ, Praekelt U, Debrand E, et al. Studies on the morphology and spreading of human endothelial cells define key inter- and intramolecular interactions for talin1. European Journal of Cell Biology. 2010;89(9):661–73.

70. Li X, Goult BT, Ballestrem C, Zacharchenko T. The structural basis of the talin-KANK1 interaction that coordinates the actin and microtubule cytoskeletons at focal adhesions. Open Biol. 2023;13(6):230058.

71. Gough RE, Jones MC, Zacharchenko T, Le S, Yu M, Jacquemet G, et al. Talin mechanosensitivity is modulated by a direct interaction with cyclin-dependent kinase-1. J Biol Chem. 2021;297(1):100837.

72. Lukas F, Duchmann M, Maritzen T. Focal adhesions, reticular adhesions, flat clathrin lattices: what divides them, what unites them? Am J Physiol Cell Physiol. 2025;328(1):C288–C302.

73. Hakanpaa L, Abouelezz A, Lenaerts AS, Culfa S, Algie M, Barlund J, et al. Reticular adhesions are assembled at flat clathrin lattices and opposed by active integrin alpha5beta1. J Cell Biol. 2023;222(8).

