## Supplementary Table S1 for "KANK2 at focal adhesion regulates their maintenance and dynamics, while at fibrillar adhesions it influences cell migration via microtubule-dependent mechanism"

### Additional file 1

N. Stojanović[1,*,#, ORCID:0000-0002-7763-4154], A. Rac[1,*, ORCID:0000-0001-8821-3059], M. Lončarić[1, ORCID:0000-0002-5343-0368], A. Tadijan[1,2, ORCID:0000-0002-5487-3611], M. Paradžik[1,3, ORCID:0000-0003-1025-5595], M. Acman[1], J.D. Humphries[4, ORCID:0000-0002-8953-7079], M.J. Humphries[5, ORCID:0000-0002-4331-6967], A. Ambriović-Ristov[1,#, ORCID:0000-0001-7784-2466]

[1]Laboratory for Cell Biology and Signalling, Division of Molecular Biology, Ruđer Bošković Institute, Zagreb, Croatia; [2]present address: Laboratory for Cell Biology, Division of Molecular Biology, Ruđer Bošković Institute, Zagreb, Croatia; [3]present address: Laboratory of Experimental Therapy, Division of Molecular Medicine, Ruđer Bošković Institute, Zagreb, Croatia, [4]Department of Life Science, Manchester Metropolitan University, Manchester, United Kingdom; [5]Manchester Cell-Matrix Centre, Faculty of Biology, Medicine & Health, University of Manchester, Manchester, United Kingdom

*equal contribution

**Table S1**

**Supplementary Table S1.** List of used antibodies and dyes.

| WESTERN BLOT | | | | | |
|---|---|---|---|---|---|
| ***Primary antibodies*** | ***Ref. No.*** | ***Distributor*** | ***Monoclonal/polyclonal*** | ***Species*** | ***Dilution*** |
| Anti-Liprin β1 | sc-514575 | Santa Cruz Biotechnology, USA | Monoclonal | Mouse | 1:100 in 5% milk |
| Anti-FAK antibody [EP695Y] | ab40794 | Abcam, USA | Monoclonal | Rabbit | 1:1000 in 5% milk |
| Anti-Paxillin [Y113] | ab32084 | Abcam, USA | Monoclonal | Rabbit | 1:7500 in 5% milk |
| Anti-EEA1 | 2411 | Cell Signaling Technology, USA | Polyclonal | Rabbit | 1:1000 in 5% milk |
| Anti-LDH | sc33781 | Santa Cruz Biotechnology, USA | Polyclonal | Rabbit | 1:400 in 5% milk |
| Anti-plectin (10F6) | sc-33649 | Santa Cruz Biotechnology, USA | Monoclonal | Mouse | 1:200 in 5% milk |
| Anti-human talin2 | MCA4771GA | Bio-Rad, USA | Monoclonal | Mouse | 1:1000 in 5% milk |
| Anti-IQGAP1 | ab133490 | Abcam, USA | Monoclonal | Rabbit | 1:1000 in 5% milk |
| Anti-vinculin | ab129002 | Abcam, USA | Monoclonal | Rabbit | 1:1000 in 5% milk |
| Anti-α-actinin 1(H-2) | sc-17829 | Santa Cruz Biotechnology, USA | Monoclonal | Mouse | 1:500 in 5% milk |
| Anti-α-actinin-4 (G-4) | sc-390205 | Santa Cruz Biotechnology, USA | Monoclonal | Mouse | 1:250 in 5% milk |
| Anti-Integrin β5 | D24A5 | Cell Signaling Technology, USA | Monoclonal | Mouse | 1:1000 in 5% milk |
| Anti-zyxin | sc-136128 | Santa Cruz Biotechnology, USA | Monoclonal | Mouse | 1:500 in 5% milk |
| Anti-KANK2 | HPA015643 | Sigma-Aldrich, USA | Polyclonal | Rabbit | 1:1000 in 5% milk |
| ***Secondary antibodies*** | ***Ref. No.*** | ***Distributor*** | ***Monoclonal/ polyclonal*** | ***Species*** | ***Dilution*** |
| Goat anti-rabbit IgG (H+L) | 31466 | Invitrogen, USA | Polyclonal | Goat | 1:5000 in 5% milk |
| Goat anti-mouse IgG (H+L) | G21040 | Invitrogen, USA | Polyclonal | Goat | 1:10 000 in 5% milk |
| **IMMUNOFLUORESCENCE** | | | | | |
| ***Primary antibodies*** | ***Ref. No.*** | ***Distributor*** | ***Monoclonal/ polyclonal*** | ***Species*** | ***Dilution*** |
| Anti-human talin1 | MCA4770GA | Bio-Rad, USA | Monoclonal | Mouse | 1:100 in 5% BSA |
| Anti-human talin2 | MCA4771GA | Bio-Rad, USA | Monoclonal | Mouse | 1:100 in 5% BSA |
| Anti-Integrin β5 | D24A5 | Cell Signaling Technology, USA | Monoclonal | Mouse | 1:600 in 5% BSA |
| Anti-KANK2 | HPA015643 | Sigma-Aldrich, USA | Polyclonal | Rabbit | 1:100 in 5% BSA |
| Anti-Integrin α5 | NBP2-50146 | Novus Biologicals, USA | Monoclonal | Mouse | 1:500 in 5% BSA |
| Anti-alpha tubulin | CP06 | Sigma-Aldrich, USA | Monoclonal | Mouse | 1:20 in 5% BSA |
| Anti-Filamin 1 (E-3) | sc-17749 | Santa Cruz Biotechnology, USA | Monoclonal | Mouse | 1:100 in 5% BSA |

| | | | | | |
|---|---|---|---|---|---|
| Anti-Filamin B | ab97457 | Abcam, USA | Polyclonal | Rabbit | 1:50 in 5% BSA |
| Anti-α-actinin 1(H-2) | sc-17829 | Santa Cruz Biotechnology, USA | Monoclonal | Mouse | 1:50 in 5% BSA |
| Anti-α-actinin-4 (G-4) | sc-390205 | Santa Cruz Biotechnology, USA | Monoclonal | Mouse | 1:50 in 5% BSA |
| Anti-vinculin | ab129002 | Abcam, USA | Monoclonal | Rabbit | 1:100 in 5% BSA |
| Recombinant Alexa Fluor® 647 Anti-Vinculin | ab196579 | Abcam, UK | Monoclonal | Rabbit | 1:200 in 5% BSA |
| ***Secondary antibodies*** | ***Ref. No.*** | ***Distributor*** | ***Monoclonal/polyclonal*** | ***Species*** | ***Dilution*** |
| Anti-Mouse IgG Alexa Fluor 546 | A-11030 | Invitrogen, USA | Polyclonal | Goat | 1:1000 in 5% BSA |
| Anti-Mouse IgG Alexa Fluor 488 | #4408 | Cell Signaling Technology, USA | | Goat | 1:1000 in 5% BSA |
| Anti-Mouse IgG Alexa Fluor 405 | A-31553 | Invitrogen, USA | Polyclonal | Goat | 1:250 in 5% BSA |
| Anti-Rabbit IgG Alexa Flour, 555 | A-31572 | Invitrogen, USA | Polyclonal | Donkey | 1:1000 in 5% BSA |
| Anti-Rabbit IgG Alexa Fluor 647 | #4414 | Cell Signaling Technology, USA | Polyclonal | Goat | 1:1000 in 5% BSA |
| Anti-Mouse IgG1 Alexa Fluor 555 | A-21127 | Invitrogen, USA | Polyclonal | Goat | 1:1000 in 5% BSA |
| Anti-Mouse $IgG2_b$ Alexa Fluor 488 | A-21141 | Invitrogen, USA | Polyclonal | Goat | 1:1000 in 5% BSA |
| ***Dyes*** | ***Ref. No.*** | ***Distributor*** | | | ***Dilution*** |
| Phalloidin, Alexa Fluor 488 | P5282 | Sigma Aldrich, USA | | | 1:100 in 5% BSA |
| **PROXIMITY LIGATION ASSAY (PLA)** | | | | | |
| ***Primary antibodies*** | ***Ref. No.*** | ***Distributor*** | ***Monoclonal/polyclonal*** | ***Species*** | ***Dilution*** |
| Anti-Integrin β5 | D24A5 | Cell Signaling Technology, USA | Monoclonal | Mouse | 1:4000 in diluent* |
| Anti-human talin2 | MCA4771GA | Bio-Rad, USA | Monoclonal | Mouse | 1:4000 in diluent* |
| Anti-human talin1 | MCA4770GA | Bio-Rad, USA | Monoclonal | Mouse | 1:4000 in diluent* |
| Anti-KANK2 | HPA015643 | Sigma-Aldrich, USA | Polyclonal | Rabbit | 1:4000 in diluent* |
| Anti-Integrin α5 | NBP2-50146 | Novus Biologicals, USA | Monoclonal | Mouse | 1:4000 in diluent* |
| ***Secondary antibodies*** | ***Ref. No.*** | ***Distributor*** | ***Monoclonal/ polyclonal*** | ***Species*** | ***Dilution*** |
| Navenibody M1 (40X) | NB.1.100.06 | Navinci Diagnostics AB, Sweden | | Mouse | 1:40 in diluent* |
| Navenibody R2 (40X) | NB.1.100.07 | Navinci Diagnostics AB, Sweden | | Rabbit | 1:40 in diluent* |

** part of NaveniFlexTM Cell MR kit*
