## Supplementary Figures S1-S9 for "KANK2 at focal adhesion regulates their maintenance and dynamics, while at fibrillar adhesions it influences cell migration via microtubule-dependent mechanism"

### Additional file 2

N. Stojanović[1,*,#, ORCID:0000-0002-7763-4154], A. Rac[1,*, ORCID:0000-0001-8821-3059], M. Lončarić[1, ORCID:0000-0002-5343-0368], A. Tadijan[1,2, ORCID:0000-0002-5487-3611], M. Paradžik[1,3, ORCID:0000-0003-1025-5595], M. Acman[1], J.D. Humphries[4, ORCID:0000-0002-8953-7079], M.J. Humphries[5, ORCID:0000-0002-4331-6967], A. Ambriović-Ristov[1,#, ORCID:0000-0001-7784-2466]

[1]Laboratory for Cell Biology and Signalling, Division of Molecular Biology, Ruđer Bošković Institute, Zagreb, Croatia; [2]present address: Laboratory for Cell Biology, Division of Molecular Biology, Ruđer Bošković Institute, Zagreb, Croatia; [3]present address: Laboratory of Experimental Therapy, Division of Molecular Medicine, Ruđer Bošković Institute, Zagreb, Croatia, [4]Department of Life Science, Manchester Metropolitan University, Manchester, United Kingdom; [5]Manchester Cell-Matrix Centre, Faculty of Biology, Medicine & Health, University of Manchester, Manchester, United Kingdom

*equal contribution

### Supplementary Methods

#### SDS-PAGE and Western Blotting (related to Supplementary Fig. S1 and S2)

Total cell lysates were obtained from 3.5 cm Petri dishes in 200 µL RIPA buffer supplemented with protease inhibitor cocktail (ThermoFisher). Samples for SDS-PAGE were collected by scraping. Samples containing an equal amount of protein were mixed in 6× Laemmli loading buffer (375 mM Tris-HCl (pH 6.8), 30% (w/v) glycerol, 12% (w/v) SDS, 0.02% (w/v) bromophenol blue, 12% (v/v) 2-mercaptoethanol) to reach a final 1× concentration, sonicated and heated for 5 min at 96°C. Isolated IACs were prepared for SDS-PAGE by solubilization in 2× Laemmli loading buffer and heating for 20 min at 70°C while shaking (1000 rpm). All samples were loaded onto pre-casted gradient gel (4 – 15% Mini-PROTEAN TGX) (Bio-Rad), separated by SDS-PAGE and semi-dry transferred to nitrocellulose (Bio-Rad). The membrane was blocked in 5% (w/v) non-fat dry milk or 5% (w/v) bovine serum albumin (BSA, Carl Roth), and incubated overnight with the appropriate antibodies, followed by incubation with horseradish peroxidase coupled secondary antibody. The primary and secondary antibodies are listed in Additional file 8: Table S1. Detection was performed using chemiluminescence (PerkinElmer) and documented with Uvitec Alliance Q9 mini (BioSPX b.v.). Blots were quantified using ImageJ.

**Supplementary Figures**

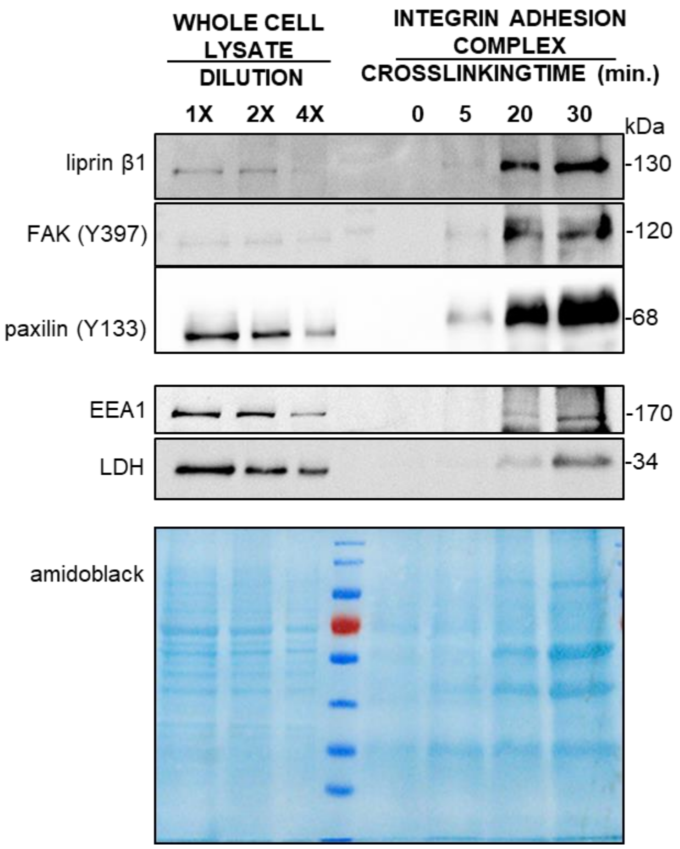


**Supplementary Fig. S1** Optimization of crosslinking duration in RPMI-7951 cells. The optimal crosslinking duration of 15 minutes was selected based on the WB analysis of marker IAC components, liprin β1, FAK (Y397) and paxillin (Y133). WB analysis of components that do not classically compartmentalize within adhesion complexes, early endosome antigen 1 (EEA1) and lactate dehydrogenase (LDH), was used to estimate co-purifying contaminants. Whole cell lysates were used as positive control. Amidoblack staining of the membrane was used as a loading control.

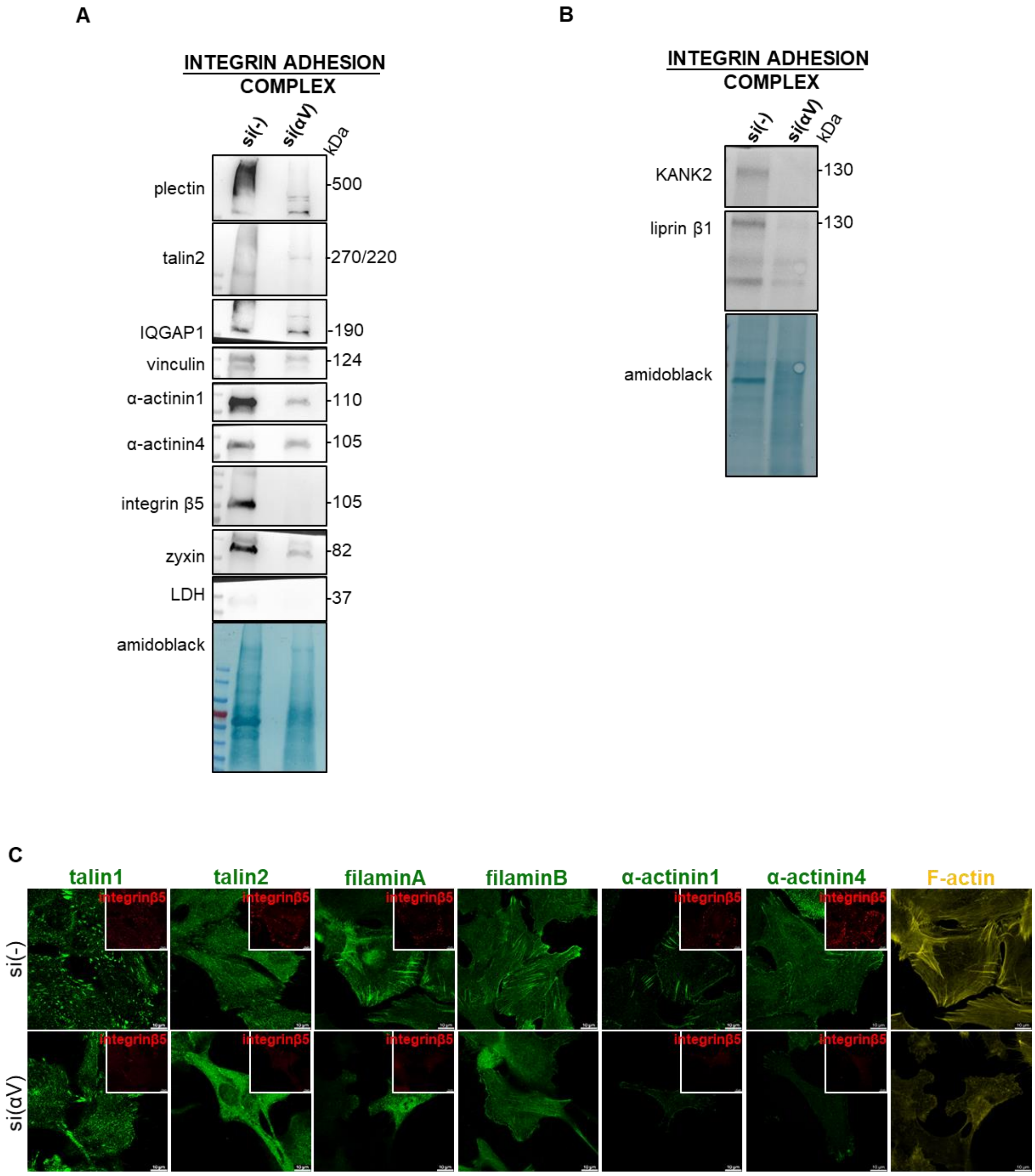


**Supplementary Fig. S2** MS data validation in RPMI-7951 cells confirms that integrin αV knockdown leads to reduced expression of several IAC proteins. (A, B) WB analysis of IAC proteins in RPMI-7951 cells transfected with either non-specific siRNA (si(-)) or integrin αV specific siRNA (si(ITGAV)). Forty-eight hours upon transfection, IACs were isolated and WB analysis was performed. Amidoblack staining of the membrane was used as a loading control. The results presented are representative of two independent experiments yielding similar results. (C) Analysis of localization of IAC proteins in RPMI-7951 cells transfected with either non-specific siRNA (si(-)) or integrin αV specific siRNA (si(ITGAV)) using immunofluorescence. Forty-eight hours upon transfection, RPMI-795 cells were fixed, permeabilized, incubated with antibodies against vinculin, talin1, talin2, filaminA, filaminB, a-actinin 1, a-actinin 4 antibody, followed by Alexa-Fluor 488-conjugated antibody (green). Where possible additional incubation with integrin β5 following incubation with Alexa-Fluor 555-conjugated antibody (shown in red) was performed. F-actin staining (shown in yellow) was performed. Analysis was performed using TCS SP8 Leica. Scale bar = 10 μm.

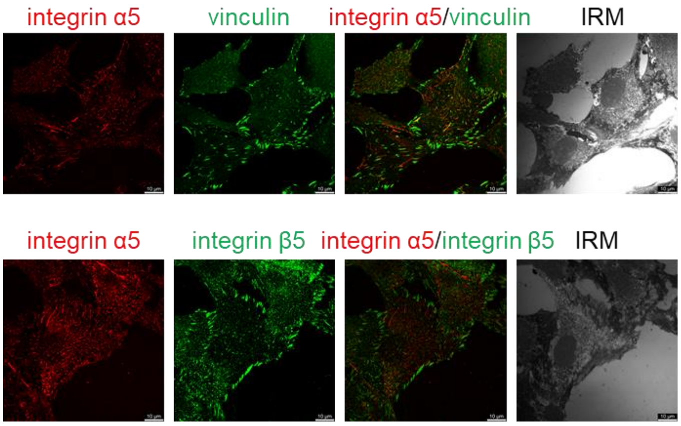


**Supplementary Fig. S3** RPMI-7951 cells contain α5-positive adhesions. Forty-eight hours upon seeding on coverslips, cells were fixed with PFA, permeabilised and stained with anti-integrin α5 and anti-vinculin, or anti-integrin β5 antibody, followed by Alexa-Fluor 546-conjugated antibody (red) or Alexa-Fluor 488-conjugated antibody (green), respectively. IRM images were taken. Analysis was performed using TCS SP8 Leica. Scale bar = 10 μm.

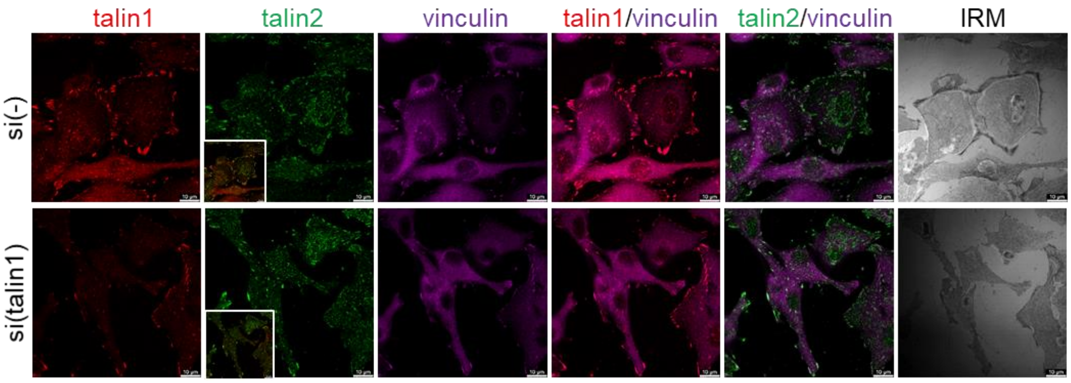


**Supplementary Fig. S4** In RPMI-7951 cells talin2 is part of talin1- and vinculin-negative structures. Forty-eight hours upon transfection with either non-specific siRNA (si(-)) or talin1 specific siRNA (si(talin1)), cells were fixed with methanol, permeabilised and stained with anti-talin1 and anti-talin2 primary antibody, followed by Alexa-Fluor 546-conjugated antibody (red) or Alexa-Fluor 488-conjugated antibody (green). Vinculin was visualized using conjugated anti-vinculin Alexa Fluor 647 antibody (magenta) and IRM images were taken. Analysis was performed using TCS SP8 Leica. Scale bar = 10 µm.

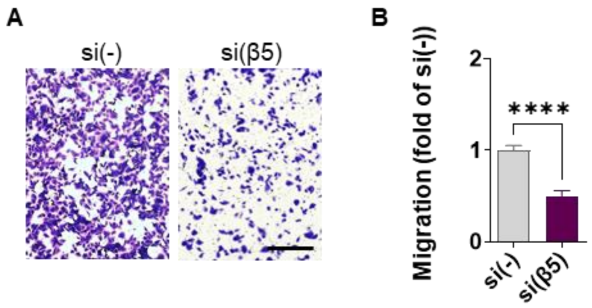


**Supplementary Fig. S5** Integrin β5 knockdown decreases migration in RPMI-7951 cells. (A) Serum starved (24 hours) cells, transfected previously with either control or integrin β5-specific siRNA, were seeded in Transwell cell culture inserts and left to migrate for 22 hours toward serum. Cells on the insert underside were stained with crystal violet, photographed, and counted. Scale bar = 100 µm. (B) Histogram data represents averages of five microscope fields of three independently performed experiments, plotted as mean ± SD. Data were analysed by unpaired Student's t-test. ns, not significant; *$P < 0.05$; **$P < 0.01$; ***$P < 0.001$; ****$P < 0.0001$.

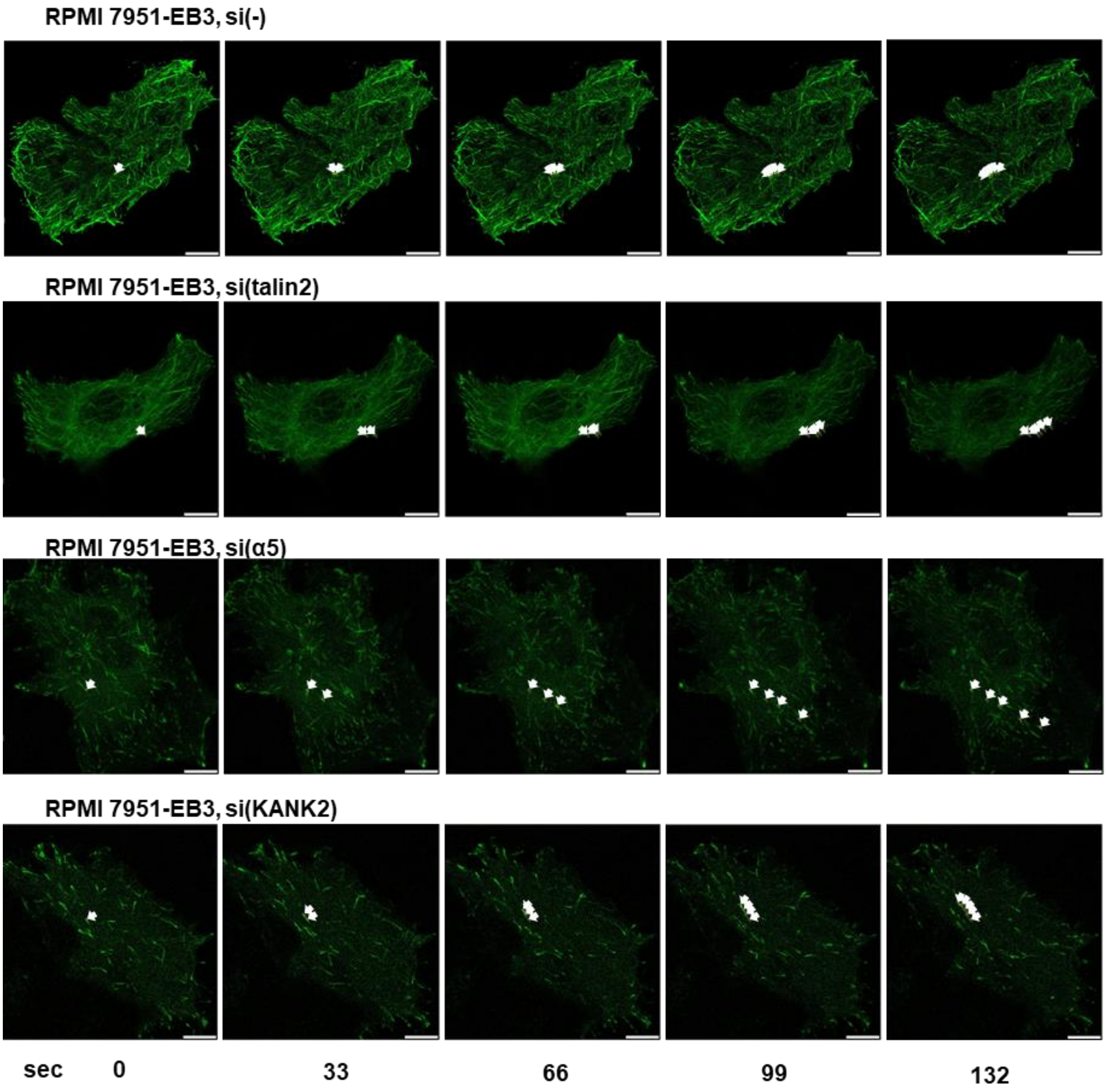


**Supplementary Fig. S6** Still images of Additional file 4: Movie S1, Additional file 5: Movie S2, Additional file 6: Movie S3 and Additional file 7: Movie S4. Images were obtained using Image J manual tracking tool. Each arrow represent position of one microtubule tip through 132 s (five frames). Images were captured every 33 s.

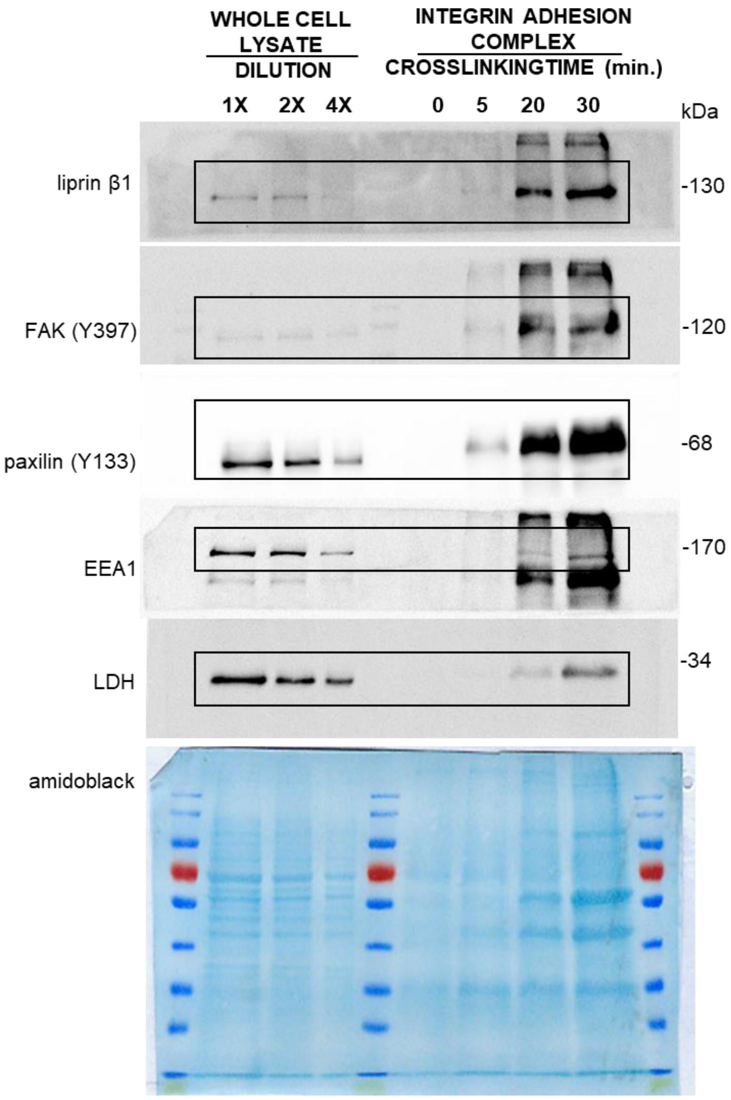


**Supplementary Fig. S7** Full images of the blots in Fig. S1. Images were obtained using Uvitec Alliance Q9 mini, which directly scanned membranes developed with ECL reagents.

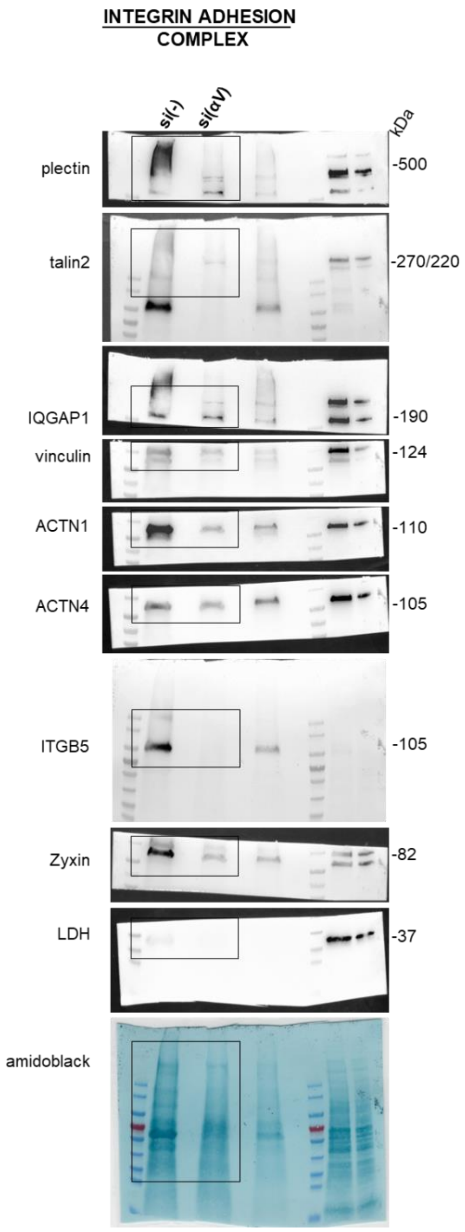


**Supplementary Fig. S8** Full images of the blots in Fig. S2A. Images were obtained using Uvitec Alliance Q9 mini, which directly scanned membranes developed with ECL reagents.

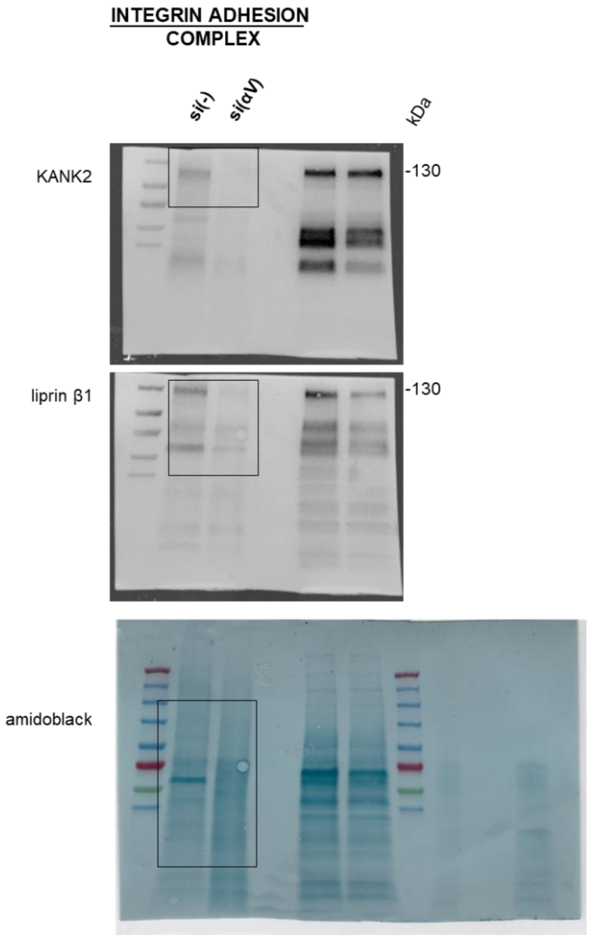


**Supplementary Fig. S9** Full images of the blots in Fig. S2B. Images were obtained using Uvitec Alliance Q9 mini, which directly scanned membranes developed with ECL reagents.

**Additional file 4: Supplementary Movie S1.** Time-lapse live cell microscopy of RPMI-7951 cells with fluorescent EB3 (RPMI-7951-EB3 cells) transfected with control siRNA used for measurement of velocity of MT growth.

**Additional file 5: Supplementary Movie S2.** Time-lapse live cell microscopy of RPMI-7951 cells with fluorescent EB3 (RPMI-7951-EB3 cells) transfected with talin2-specific siRNA used for measurement of velocity of MT growth.

**Additional file 6: Supplementary Movie S3.** Time-lapse live cell microscopy of RPMI-7951 cells with fluorescent EB3 (RPMI-7951-EB3 cells) transfected with integrin α5-specific siRNA used for measurement of velocity of MT growth.

**Additional file 7: Supplementary Movie S4.** Time-lapse live cell microscopy of RPMI-7951 cells with fluorescent EB3 (RPMI-7951-EB3 cells) transfected with KANK2-specific siRNA used for measurement of velocity of MT growth.
